# PHLDA2 promotes breast cancer metastasis by co-opting a developmental program for placental vascular remodeling

**DOI:** 10.1101/2025.08.19.670946

**Authors:** Paige V. Halas, Hannah Savage, Stephanie J. Hachey, Tatyana Lev, Angela Lincy Prem Antony Samy, Connie Chan, Stephanie Jimenez, Radhika M. Domadia, Iana Lin, Sharmila Mallya, Jacob Insua-Rodriguez, Jessica Gonzalez, Isam Adam, Hamad Alshetaiwi, Grace A. Hernandez, Aaron Longworth, Timothy P. McMullen, Kai Kessenbrock, Christopher C.W. Hughes, Dennis Ma, Devon A. Lawson

## Abstract

Identifying drivers of metastasis is essential for developing new treatments for patients with advanced disease. Here, we identify *PHLDA2* as a robust driver of breast cancer metastasis. Previous work established *PHLDA2* as an imprinted gene expressed by trophoblasts which are critical for vascular remodeling during placental development. We find that hypomethylation of *PHLDA2* in breast tumors correlates with increased gene expression, which is associated with metastasis and poor survival in breast cancer patients. RNA-sequencing showed that *PHLDA2* overexpression results in upregulation of genes that control invasion, extracellular matrix assembly, and vascular remodeling, consistent with trophoblast functions in placental development. Using an *in vitro* vascularized microtumor (VMT) system, we find that *PHLDA2* functions through *SPARC,* which promotes metastasis by inducing vascular permeability and enhancing tumor dissemination. These data suggest that increased expression of *PHLDA2* through hypomethylation promotes metastasis by ectopic expression of a developmental program for vascular remodeling.

## Introduction

Cancer metastasis, the primary cause of cancer-related mortality, is increasingly linked to the aberrant reactivation of latent developmental programs normally quiescent in adult tissues. Cancer cells can hijack developmental pathways governing processes such as self-renewal, angiogenesis and epithelial-mesenchymal transition (EMT) to gain migratory, invasive, and survival advantages^1,2^. This reinstatement of embryonic programs is often orchestrated by epigenetic reprogramming that can unlock silenced developmental gene networks^3^. This grants cancer cells the plasticity needed to navigate the multi-step metastatic cascade in which a cancer cell leaves the primary tumor, travels through the bloodstream, and establishes metastatic lesions at a distant site^4^. The development of new treatments that target developmental pathways hijacked by cancer cells therefore offers a promising strategy to prevent metastatic spread.

In previous work, we screened for cellular programs important for metastasis using single-cell RNA sequencing (scRNAseq)^5,6^. This identified pleckstrin homology-like domain family A member 2 (*PHLDA2*) as a top gene upregulated in metastatic cells^5^. *PHLDA2* was first named Imprinted in Placenta and Liver (*IPL*) in the late 1990s when it was identified as a maternally expressed imprinted gene tightly regulated by epigenetic methylation^7^. The function of *PHLDA2* has been studied most extensively in placental development where its overexpression due to hypomethylation and re-expression of the paternal allele results in reduced placental growth and fetal growth restriction in both mice and humans^8–12^. The placenta is a highly vascularized organ that supports fetal development by facilitating the exchange of nutrients, oxygen, and waste products between the mother and fetus^13,14^. *Phlda2* is primarily expressed by trophoblasts that are derived from the blastocyst and invade into the maternal endometrium to generate the placenta following implantation^15^. They subsequently diversify, secreting factors that induce vascular remodeling as well as generating key structural components of the feto-maternal interface^13,14^. Studies in genetically engineered mouse models have shown that inappropriate expression of *Phlda2* results in aberrant trophoblast differentiation, placental development and nutrient exchange^8–11,16^.

Interestingly, *PHLDA2* also plays important roles in cancer. Initial reports suggested a tumor-suppressive function in osteosarcoma, lymphoma, and lung cancer^17,18^. Recent work has also shown a tumor promoting role, where *PHLDA2* promotes cancer progression through Akt pathway activation^19,20^, angiogenesis^21^, and enhancement of cancer cell stemness^21^. Like in placental development, altered *PHLDA2* expression in cancer may occur through dysregulated imprinting^22,23^. In clear cell renal carcinoma, hypomethylation of the *PHLDA2* locus correlates with overexpression and predicts poor therapeutic response^24^. In breast cancer, increased *PHLDA2* expression has been associated with poor prognosis^25^, but its function and role in disease progression remain unclear.

Here, we find that *PHLDA2* is a robust driver of breast cancer metastasis through vascular remodeling. Analysis of published datasets shows that increased *PHLDA2* expression is associated with increased metastasis and reduced survival in breast cancer patients. Analysis of TCGA methylation microarray data revealed that decreased methylation at two specific CpG sites located in the untranslated exon of *PHLDA2* is associated with increased gene expression, suggesting that *PHLDA2* overexpression in breast tumors occurs through hypomethylation and loss of imprinting. In mouse models, we find that lentivirally-mediated overexpression of *PHLDA2* results in a robust increase in lung metastasis, showing that *PHLDA2* functionally promotes metastasis. RNA-sequencing showed that *PHLDA2* overexpression is associated with upregulation of transcriptional programs that regulate invasion, extracellular matrix assembly, and vascular remodeling, remarkably consistent with trophoblast functions in placental development. Using an *in vitro* vascularized microtumor (VMT) system^26–28^, we found that *PHLDA2* overexpression induces vascular permeability, which is understood to promote metastasis by enhancing tumor cell intravasation and extravasation^29^. We further show that it exerts its function through upregulation of Secreted Protein Acidic and Cysteine Rich *(SPARC)*, which has been shown to promote metastasis through vascular permeability^30^. These data support a model where increased expression of *PHLDA2* through hypomethylation promotes breast cancer metastasis by ectopic expression of a developmental program for vascular remodeling that is characteristic of placental trophoblasts.

## Results

### Spatiotemporal expression of *Phlda2* in the developing murine placenta

The placenta is a transient, yet vital organ that forms the primary link between the mother and fetus. This is due to its hemochorial structure characterized by direct contact between maternal blood and fetal trophoblasts. In the murine placenta, chorioallantoic attachment, where the fetal allantois fuses with the chorionic plate, initiates the development of the two main functional layers in the placenta: the labyrinth and the junctional zone which both contain multiple trophoblast populations^14,31^ (**Figure 1A**). The labyrinth, containing a vascular network that serves as the primary site for feto-maternal exchange, lies adjacent to the junctional zone, which arises from the ectoplacental cone^14,31^ (**Figure 1A**). The junctional zone is situated next to the decidua—the modified maternal uterine lining that the placenta invades. The placenta is encased by the outer muscular layer of the uterus, or the myometrium, and in a murine model reaches full structural maturity by E14.5 after successfully remodeling maternal arteries to ensure blood flow^14,31^ (**Figure 1A**). *Phlda2* is critical for the formation of these placental zones in addition to trophoblast function^8,9^. Loss of *Phlda2* leads to an expanded spongiotrophoblast and glycogen trophoblast layer in the junctional zone^8^, suggesting it plays complex, context-dependent roles in multiple trophoblast lineages.

**Figure 1:**
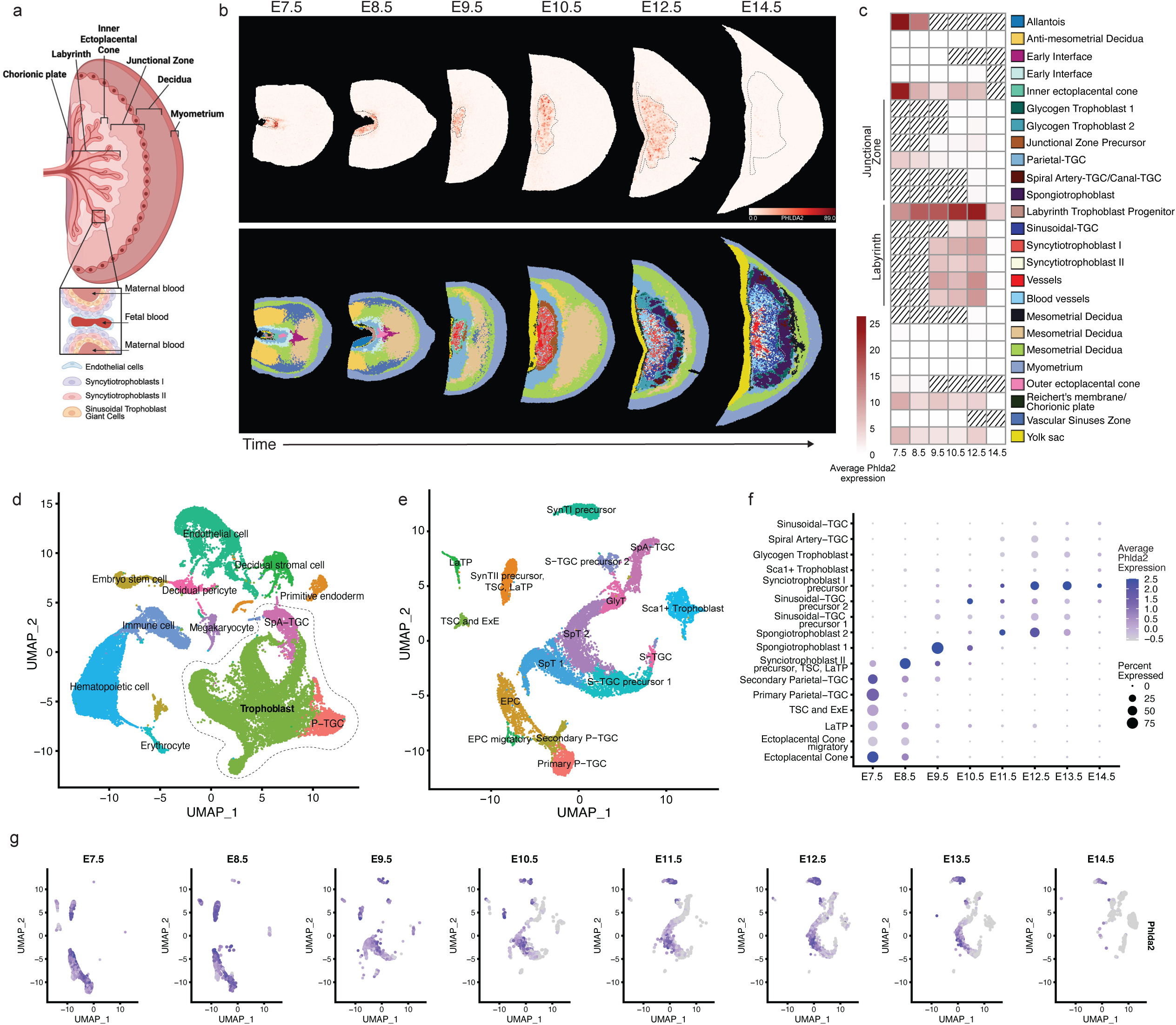
Spatiotemporal expression of *Phlda2* in the developing placenta. (a) Schematic of the murine placenta highlighting vascular architecture and trophoblast populations involved in vascular development. The labyrinth zone, where maternal and fetal blood exchange occurs, contains a dense network of fetal capillaries surrounded by trophoblast subtypes. Syncytiotrophoblasts (SynT-I and SynT-II) and sinusoidal trophoblast giant cells (S-TGC) are shown spatially interacting with fetal blood and play key roles in regulating vascular patterning, remodeling, and barrier function. (b) Top: Spatial mapping of *Phlda2* transcripts (red) in tissue sections of developing placenta from E7.5-14.5. Bottom: Schematic representing spatial location of placental cell types present at each stage (color key in (c)). Black or white dotted lines denotes the allantois, chorionic plate, and/or labyrinth area in each tissue section which the majority of *Phlda2* is localized within. (c) Heatmap depicting *Phlda2* expression levels across placental cell types and zones across developmental time. Color key corresponds to (b). (d) UMAP projection of single cell RNA sequencing data of mouse placenta. Clusters represent placental cell populations. Trophoblast populations are highlighted with a black dotted line. (e) UMAP projection of trophoblasts subset (highlight in (d)) from single cell RNA sequencing data of mouse placenta, showing distinct trophoblast cell populations and their transcriptional heterogeneity. (f) Dot plot displaying *Phlda2* gene expression levels in trophoblast cell populations across developmental timepoints. Color gradient represents average *Phlda2* expression and dot size represents the precent of the population expressing *Phlda2*. (g) UMAP projections of trophoblasts showing *Phlda2* expression at each developmental time.

In order to gain insights into the functional role of *Phlda2*, we first aimed to temporally and spatially map gene expression of trophoblast populations during placental development. To investigate this, we analyzed the spatial distribution and transcriptomic expression of *Phlda2* across murine placental development using publicly available datasets. We first analyzed a murine spatial transcriptomics dataset comprising of six embryonic tissues from E7.5 to E14.5^32^. Visualizing *Phlda2* transcripts revealed dynamic expression patterns (**Figure 1B**), which we quantified across the cell types annotated in the dataset (**Figure 1C**). At early stages (E7.5-E8.5), high *Phlda2* expression was prominent in precursor structures, including the allantois, chorionic plate, and inner ectoplacental cone (EPC), which is a known reservoir for trophoblast stem cells (TSC)^33^. Following chorioallantoic fusion around E8.5, as the labyrinth begins to form^14^, *Phlda2* expression becomes highly enriched in Labyrinth Trophoblast Progenitor (LaTP) cells, peaking between E8.5 and E12.5. LaTPs are essential for generating the specialized trophoblast subtypes of the labyrinth, including Syncytiotrophoblast I and II (SynT I/SynT II) and Sinusoidal-Trophoblast Giant Cells (S-TGC)^34^, which also show *Phlda2* expression from E9.5 to E12.5 as they form the critical feto-maternal exchange interface. By E14.5, as the placenta reaches structural maturity, *Phlda2* expression subsides in these labyrinthine trophoblast populations.

To overcome the resolution limits of the spot-based spatial analysis and precisely define trophoblast subpopulations that express *Phlda2*, we additionally leveraged a scRNAseq dataset of mouse placental development containing tissues from eight timepoints spanning from E7.5 to E14.5^35^ (**Figure 1D**). Confirming our initial findings, *Phlda2* expression was largely restricted to the trophoblast lineage (**Figure S1A-B**) prompting us to subset these cells for more granular annotation (**Figure 1E, see Methods**). Re-clustering revealed the presence of trophoblast subpopulations not defined in the spatial data including Primary and Secondary Parietal-Trophoblast Giant Cells (P-TGC) and multiple trophoblast precursor populations (**Figure 1E**). Analysis of the *Phlda2* expression over time revealed a dynamic pattern that mirrored our spatial analysis (**Figure 1F-G, S1C**). At the earliest timepoint (E7.5), *Phlda2* was highly expressed in stem cell and progenitor populations, including the EPC, TSCs, LaTPs, and early differentiating Primary and Secondary P-TGCs (**Figure 1F-G**). As development proceeded through mid-gestation (E9.5-E12.5) high *Phlda2* expression was maintained in labyrinth trophoblast populations, most notably the precursors of S-TGC, SynT I and SynT II, which together form the essential feto-maternal exchange barrier^14^ (**Figure 1A**). Consistent with our spatial analysis, as the placenta matures from E13.5 and E14.5 *Phlda2* expression decreased across trophoblast subtypes, suggesting its principal role is in regulating the proliferation and fate decisions of early placental progenitors rather than maintaining the function of mature trophoblasts. Together, our analysis reveals that *Phlda2* expression is highly dynamic and expressed across virtually all trophoblast subpopulations during placental development. However, it is particularly enriched in labyrinth progenitor populations and their daughter cell lineages indicating that *Phlda2* may be critical for the function of trophoblasts involved in vessel remodeling and nutrient exchange in the placenta.

### *PHLDA2* expression is associated with increased metastasis and poor survival in breast cancer patients

To evaluate the relevance of *PHLDA2* in human breast cancer patients, we analyzed its expression in healthy breast tissues (n=114) and invasive breast carcinoma tissues (n=1097) from The Cancer Genome Atlas (TCGA) dataset^36,37^. This revealed that *PHLDA2* expression is 3 to 16-fold higher in patient breast tumors relative to healthy breast tissue (**Figure 2A**). We further found that *PHLDA2* shows the highest expression in the HER2+ breast cancer subtype (**Figure S2A**), consistent with prior reports that *PHLDA2* is regulated by HER2 signaling^19^. Next, we evaluated whether *PHLDA2* expression predicts survival outcomes in breast cancer patients. This demonstrated that high *PHLDA2* expression in primary breast tumors is associated with reduced relapse free survival (RFS, n=4929) and distant metastasis free survival (DMFS, n=2765) in breast cancer patients^38,39^ (**Figure 2B-C**). High *PHLDA2* expression remained associated with poor RFS in nearly all subtypes when patients were categorized by breast cancer subtype (**Figure S2B-F**), indicating that high *PHLDA2* expression in primary tumors is a robust predictor of poor survival across breast cancer patient populations.

**Figure 2:**
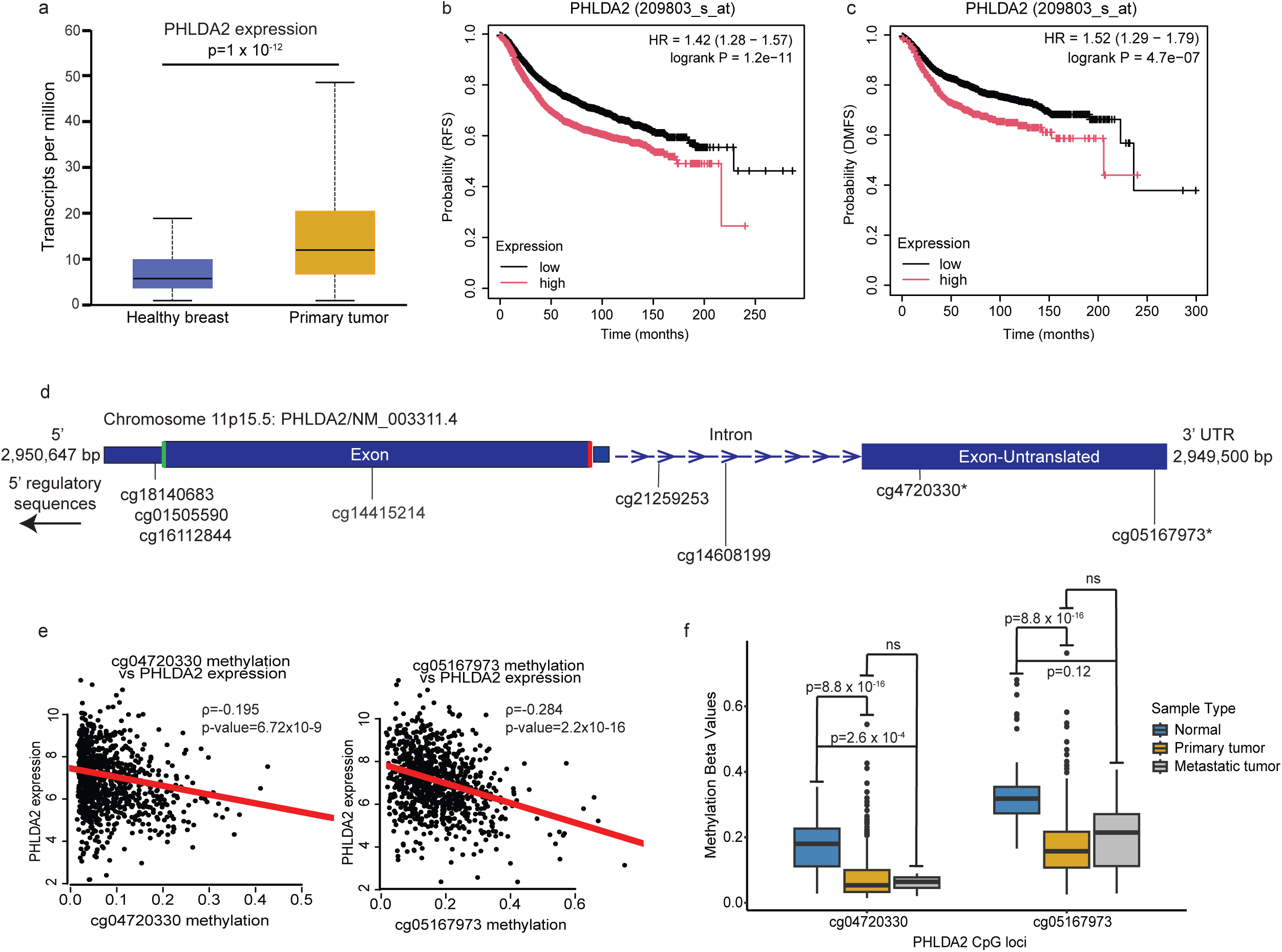
Elevated *PHLDA2* expression in patient breast tumors is associated with hypomethylation, increased metastasis and poor and hypomethylation at specific CpG loci survival. (a) *PHLDA2* expression in healthy breast (n=114) and primary breast tumor (n = 1097) tissues. Bar plot shows RNA expression in tissues from the TCGA dataset plotted using the UALCAN tool (ualcan.path.uab.edu). Data are displayed as mean ± SD. *P* value was calculated using a student’s t-test. (b) Kaplan-Meier plot showing probability of relapse free survival (RFS) in breast cancer patients with high (n=2208) vs. low (n=2721) expression of *PHLDA2* in their primary tumor tissue. Plot was generated using the KM plotter compiled dataset of mRNA breast cancer gene chip and visualized using the KM plotter tool (KMplot.com). (c) Kaplan-Meier plot showing probability of distant metastasis free survival (DMFS) in breast cancer patients with high (n=717) vs. low (n=2048) expression of *PHLDA2* in their primary tumor tissue. Plot was generated using the KM plotter compiled dataset of mRNA Breast Cancer gene chip and visualized using the KM plotter tool (KMplot.com). (d) Schematic shows CpG loci in the exonic and intronic regions of the *PHLDA2* gene. Loci located in upstream 5’ regulatory sequences are not shown. Loci were annotated using the GRCh37/hg19 human genome and the Infinium HumanMethylation450 assay. Asterisks depict hypomethylated loci in primary and metastatic tumor tissues relative to healthy breast tissues. (e) Correlation plots show relationship between *PHLDA2* RNA expression and methylation at select CpG loci (cg4720330, cg05167973). *P* value was calculated using a student’s t-test with Bonferroni multiple testing adjustment. The rho value was calculated using Spearman’s rank correlation. (f) Bar plot shows methylation values at select CpG loci (cg4720330, cg05167973) in healthy breast (n = 139), primary (n = 1101), and metastatic (n = 5) tumor tissues from breast cancer patients from the TCGA dataset. Select loci were identified in a screen for loci that display hypomethylation in tumor tissues and correlation with increased *PHLDA2* expression. DNA methylation levels, as provided in TCGA, are represented by beta values between zero and one (where 0=unmethylated and 1=methylated). Methylation levels at differentially methylated loci for each condition are displayed as box and whisker plots. Data are displayed as mean ± SD. *P* values were calculated using student’s t test with Bonferroni multiple-testing adjustment.

### *PHLDA2* hypomethylation is associated with increased gene expression in breast cancer patients

*PHLDA2* was initially characterized as an imprinted gene^7^ therefore we sought to determine whether epigenetic mechanisms similarly regulate *PHLDA2* in tumors. *PHLDA2* is a 1,148 kb gene on chromosome 11 that is composed of two exons (one untranslated), one intron, and 5’ upstream regulatory sequences that include the promoter region. There are 32 CpG loci known to be associated with *PHLDA2*, including six exonic, two intronic, and 24 in the 5’ regulatory sequence (**Figure 2D**). We calculated mean methylation levels at each locus and compared primary breast cancer tissues to healthy breast tissues using TCGA methylation microarray data^40^. We found that most loci exhibit low beta values (mean: 0.05), suggesting they remain unmethylated in both healthy breast and tumor tissues (**Figure S2G**). However, two loci located in the untranslated exon region (cg04720330 and cg051679730) show higher beta values (cg04720330 normal median: 0.18, and cg05167973 normal median: 0.32) and appear methylated in healthy breast tissue (**Figure S2G**). Importantly, we found an inverse correlation between *PHLDA2* RNA expression and methylation at these two specific loci (**Figure 2E**), suggesting that methylation at these sites may regulate *PHLDA2* expression. These loci also display lower methylation in primary and metastatic tumor samples compared to healthy tissue (**Figure 2F**). These findings suggest that the increased *PHLDA2* expression observed in primary breast and metastatic tumor samples may occur through hypomethylation or a reversal of imprinting, particularly at two loci located in the untranslated exon of the gene.

### *PHLDA2* promotes breast cancer metastasis *in vivo*

We next asked whether *PHLDA2* functionally promotes metastasis by evaluating how its genetic modulation affects breast cancer metastasis *in vivo*. We utilized a human patient-derived xenograft (PDX) model of triple negative breast cancer (HCI010) that spontaneously metastasizes to the lung following orthotopic transplantation. We previously found that cells from this model display increased *PHLDA2* expression (2-3-fold) during metastatic seeding in the lung^5^. HCI010 cells were propagated using a 3D tumor sphere culture system, followed by lentiviral infection using constructs to overexpress *PHLDA2* and GFP (*PHLDA2*-GFP) or only GFP as a control (GFP)^41^. Western blotting confirmed successful overexpression of *PHLDA2* relative to control (**Figure S3A**). Importantly, *PHLDA2* overexpression did not result in significant changes *in vitro*, indicating that the cells are viable and show similar growth kinetics after genetic modulation (**Figure S3B**).

HCI010 cells were injected orthotopically into the fourth mammary fat pad of NOD SCID gamma (NSG) mice and tissues were harvested two months later (**Figure 3A**). *PHLDA2* overexpression resulted in a limited enhancement of primary tumor growth (**Figure S3C**), demonstrating that primary tumor size should not have significant confounding effects on metastatic burden. We evaluated the impact of *PHLDA2* overexpression on lung metastasis using whole mount fluorescence microscopy and flow cytometry for GFP (**Figure 3B-C**). Remarkably, we found that *PHLDA2* overexpression leads to 6-10 times more metastatic cells in the lung than control (**Figure 3C**).

**Figure 3:**
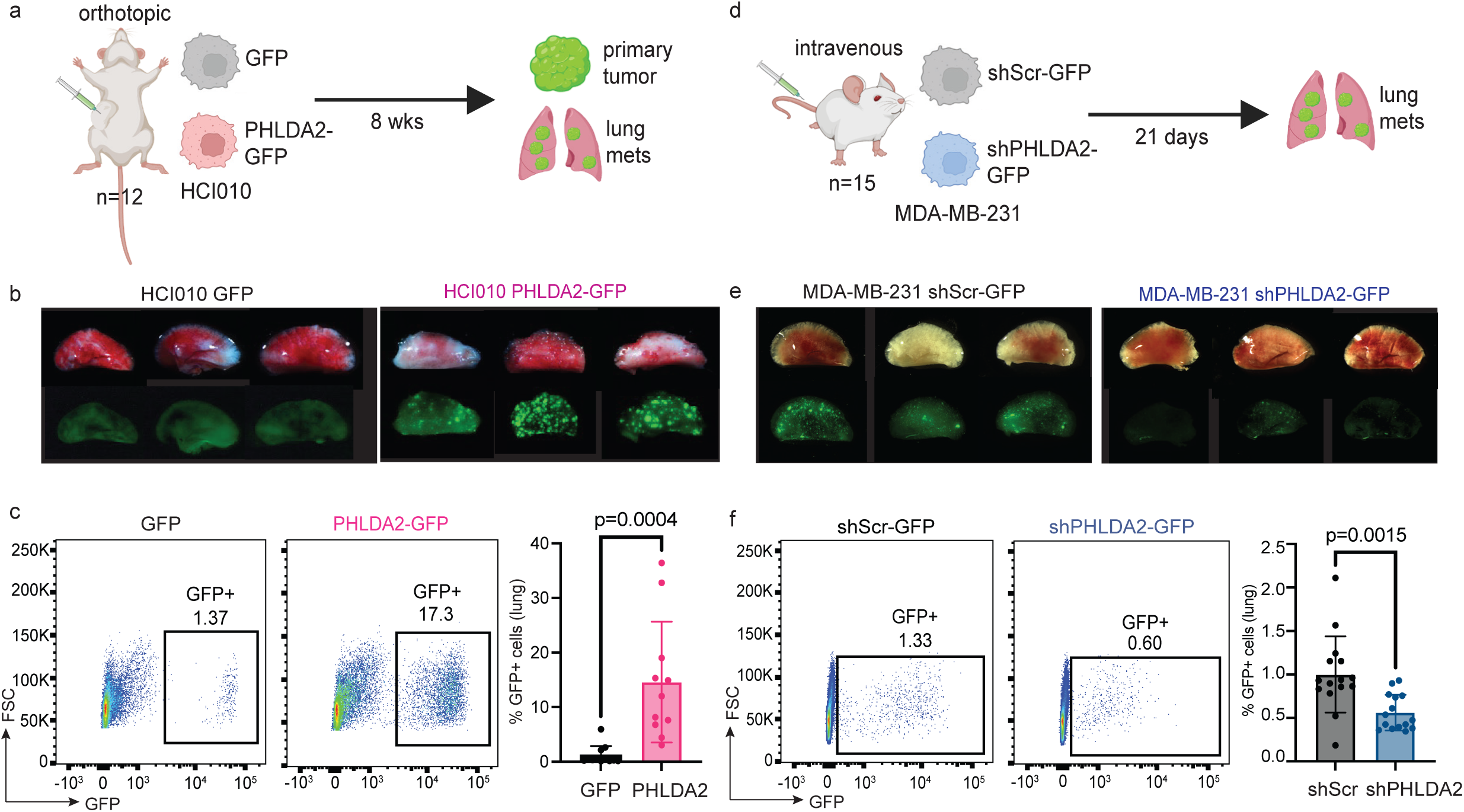
Overexpression of *PHLDA2* promotes breast cancer lung metastasis *in vivo*. (a) Schematic shows experimental design to determine how *PHLDA2* overexpression impacts spontaneous lung metastasis using the HC010 PDX model. 120,000 HCI010 tumor cells expressing GFP (control, n=12) or *PHLDA2*-GFP (n=12) were injected orthotopically into the mammary fat pad region of NSG mice. After two months, primary tumors and lungs were excised. Tumors were measured and lungs were dissociated into single cell suspensions for analysis of GFP^+^ tumor cells by flow cytometry. (b) Representative whole mount fluorescence images of lungs from transplanted mice as described in (a). (c) Representative flow cytometry (left) show quantification of GFP^+^ HCI010 cells in the lungs of transplanted mice. Bar plot (right) shows percentage of GFP^+^ HCI010 cells out of live cells in the lungs from each animal (n=16/group). Data are displayed as mean ± SD. P value was calculated using a student’s t-test and is represented on the graph. (d) Schematic shows experimental design to determine how *PHLDA2* loss impacts experimental lung metastasis using the MDA-MB-231 cell line model. 500,000 MDA-MB-231 tumor cells expressing Scrambled GFP (control) or shPHLDA2 GFP (n=15/group) were injected intravenously into NSG mice. After 21 days, lungs were excised and dissociated into single cell suspensions for analysis of GFP+ tumor cells by flow cytometry. (e) Representative whole mount fluorescence images of lungs from transplanted mice as described in (d). (f) Representative flow cytometry plots (left) show quantification of GFP^+^ MDA-MB-231 cells in the lungs of transplanted mice. Bar plot (right) shows percentage of GFP^+^ MDA-MB-231 cells out of live cells in the lung from each animal (n=15/group). Data are displayed as mean ± SD. P value was calculated using a student’s t-test and is represented on the graph.

Likely as a result of low baseline levels of *PHLDA2* expression in the HCI010 PDX model (**Figure S3D**), *PHLDA2* knockdown in the HCI010 PDX model resulted in a reduced but non-significant change in lung metastasis (**data not shown**). We therefore performed analogous experiments using MDA-MB-231 breast cancer cells, which have ∼2-fold higher levels of endogenous *PHLDA2* expression than HCI010 cells (**Figure S3D**). MDA-MB-231 cells were transduced with lentiviral constructs to knockdown *PHLDA2* (shPHLDA2-GFP) or a non-targeting Scramble as a control (shScr-GFP). Western blotting confirmed efficient knockdown of *PHLDA2* following lentiviral infection *in vitro* (**Figure S3E**). *PHLDA2* knockdown resulted in minimal changes to cell growth (**Figure S3F**). We then utilized an experimental model of metastasis by injecting the MDA-MB-231 cells via the tail vein. Lung tissues were harvested after 21 days (**Figure 3D**). Analysis of lung tissues revealed that *PHLDA2* knockdown results in a significant reduction in metastatic burden in the lung (**Figure 3E-F**). Taken together, these findings show that *PHLDA2* is not only elevated in cancer, but also functionally promotes breast cancer lung metastasis *in vivo*.

### *PHLDA2* promotes extracellular matrix and vascular remodeling gene programs

To investigate how elevated *PHLDA2* expression may drive cancer progression, we performed bulk RNA sequencing to identify associated changes in cellular programs and oncogenic pathways. Tumors were generated from HCI010 control (n=4) and PHLDA2- overexpressing (n=4) cells and dissociated to single cell suspension. Tumor cells were isolated by flow cytometry-based sorting on GFP and CD298 expression (**Figure 4A**). Libraries were prepared using the Illumina TruSeq stranded mRNA protocol and sequenced on a NovaSeq 6000 (paired-end, 100 bp, ∼50M reads/sample). Reads were aligned to the human reference genome (hg38), and differential gene expression was analyzed with DESeq2. This revealed 530 significantly upregulated and 288 downregulated genes (adjusted p value > 0.05; log2 fold change ≥ 0.5 or ≤ - 0.5 respectively) in *PHLDA2*-overexpressing tumor cells relative to controls (**Figure 4B**). Importantly, top genes were conserved across sample replicates, including *PHLDA2* confirming successful overexpression (**Figure S4A**). Gene set enrichment analysis (GSEA) revealed several recurrent cellular programs upregulated in *PHLDA2*-overexpressing tumors, including gene set terms related to extracellular matrix (ECM) organization and vascular development (**Figure 4C-E, Table 1).** Of note, we found that the top-ranked gene sets contain many overlapping genes, suggesting that *PHLDA2* may regulate a ‘meta-program’ that induces multiple pathways and promote metastasis through pleiotropic effects.

**Figure 4:**
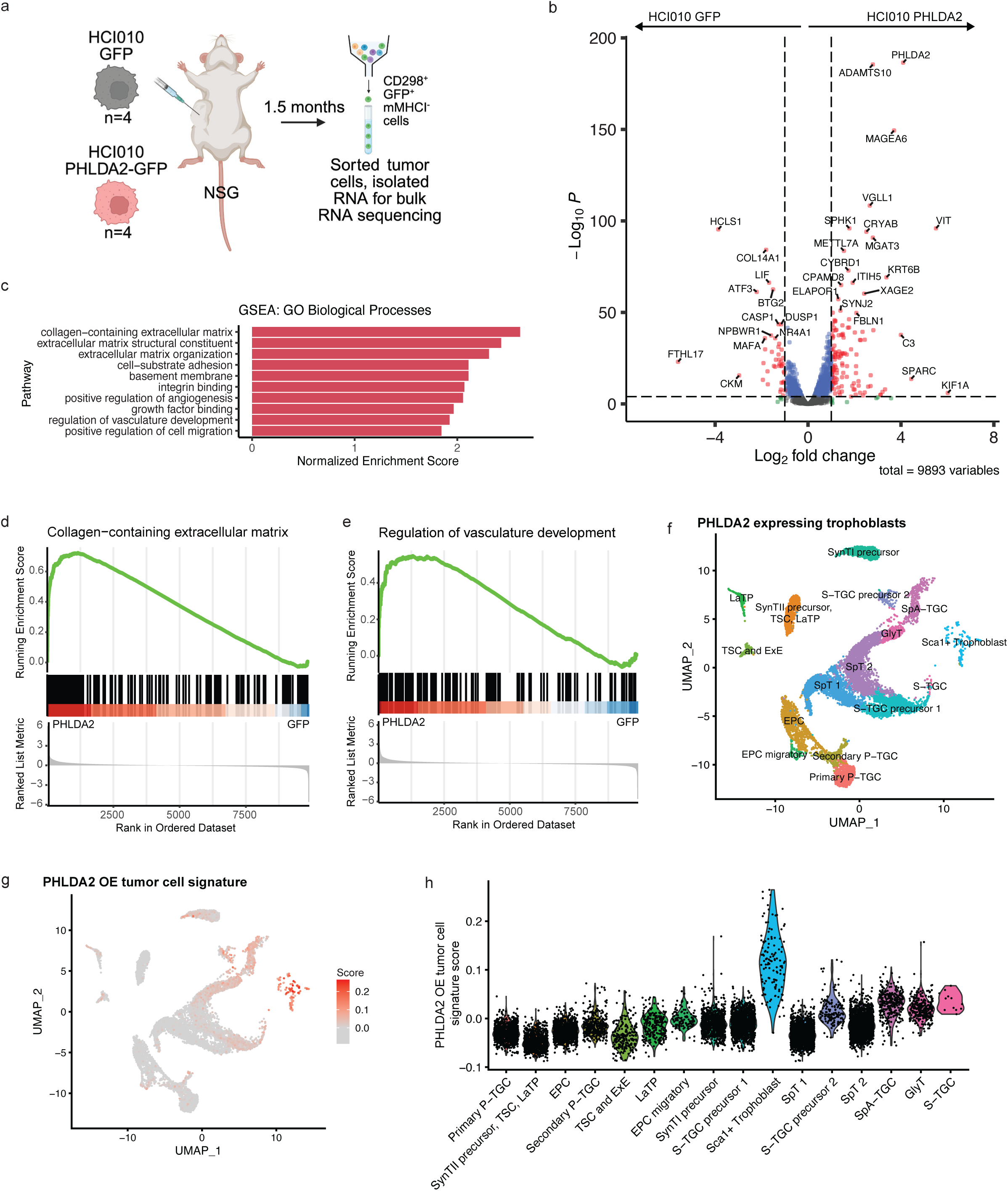
Cellular programs associated with *PHLDA2* overexpression revealed by RNA sequencing. (a) Schematic illustrates experimental design for comparison of PHLDA2-overexpressing and control tumors by RNA sequencing. HCI010 GFP and HCI010 PHLDA2 primary tumors were collected (n=4) and tumor cells were isolated by flow cytometry using GFP expression. Mouse cells were excluded using a mouse specific MHC-I antibody. RNA was isolated from tumor cells followed by library preparation and RNA sequencing. (b) Volcano plot showing differentially expressed genes in HCI010 cell overexpressing *PHLDA2* versus GFP tumor cells. Plot was generated using the R package EnhancedVolcano. (c) Bar plot depicting top ten pathways upregulated in *PHLDA2*-overexpressing tumor cells identified by gene set enrichment analysis (GSEA). GSEA was performed using R package ClusterProfiler and plot was generated with ggplot2. (d) Enrichment plot of collagen-containing extracellular matrix (GO:0062023) Gene Ontology gene set term. Plot was generated using the R package enrichplot. (e) Enrichment plot of vasculature development (GO:1901342) Gene Ontology gene set term. Plot was generated using the R package enrichplot. (f) UMAP projection of *Phlda2* expressing trophoblasts. Trophoblasts with *Phlda2* expressing > 0.1 were subset (which was 76% of total trophoblasts). (g) UMAP projection of 100 gene *PHLDA2* overexpressing tumor cell signature scoring in *Phlda2* expressing trophoblasts. (h) Violin plot of 100 gene *PHLDA2* overexpressing tumor cell signature scoring across *Phlda2* expressing trophoblasts populations.

Top-ranked genes associated with the ECM gene terms included collagen subunits (*COL1A1*, *COL1A2*, *COL2A1*), structural ECM genes (*VIT*, *MFAP5*, *LUM*), and regulators of ECM restructuring (*SPARC*). We validated several of these markers by qPCR (**Figure S4B**), and sqdew on the protein level with immunohistochemical approaches. MFAP5, a microfibril glycoprotein linked to breast cancer proliferation and migration^42^, showed significantly higher expression in metastatic lung lesions from *PHLDA2*-overexpressing tumors versus controls *in situ* (**Figure S4C-D**). We additionally quantified collagen deposition, which promotes metastasis by increasing tumor stiffness^43^, with Masson’s Trichrome staining. We found a 3- to 7-fold increase in collagen area in metastatic lesions in *PHLDA2*-overexpressing lesions (**Figure S4E-F**). These data indicate that *PHLDA2* enhances ECM deposition or remodeling.

Highly ranked genes related to vascular development and angiogenesis (**Figure 4E**) included Secreted protein acidic and rich in cysteine (*SPARC*), Complement C3 (*C3*), and Sphingosine kinase-1 (*SPHK1*). Strikingly, all are associated with vascular barrier integrity. SPARC is a secreted factor that binds to the adhesion molecule VCAM1 on activated endothelial cells^44^ causing vascular barrier disruption through a ROS-MKK3/6-p38MAPK-MLC2 signaling pathway^30^. The kinase SPHK1 catalyzes the production of the secreted factor Sphingosine-1-Phosphate (S1P), which binds to S1PR2 on endothelial cells^45^. This binding activates the RhoA/ROCK signaling pathway which reduces the translocation of translation of VE-Cadherin to adherens junctions, thereby leading to increased vascular permeability^45^. Similarly, C3 increases calcium signaling which reduces VE-cadherin expression at adherens junctions^46^.

Vascular remodeling is a critical process in labyrinth formation during placental development^14^ and recognized driver of cancer metastasis^29,47^. Thus, we hypothesized that overexpressing *PHLDA2* in breast tumor cells may increase gene programs upregulated in certain populations of trophoblasts during placental development. To test this, we generated a *PHLDA2*-overexpressing tumor cell gene signature comprising of the top 100 upregulated differentially expressed genes from our bulk RNA sequencing (**Table 2**). We utilized the previously published placenta scRNAseq dataset and subset *PHLDA2*-expressing trophoblasts (gene expression > 0.1, 76% of trophoblasts remained after subset). Within the *PHLDA2*-expressing trophoblasts, we performed gene scoring of our signature (**Figure 4F**). Several clusters had low scoring of the gene signature including the Spiral Artery trophoblast giant cells (SpA-TGC), glycogen trophoblasts (GlyT), and a Sinusoidal trophoblast giant cells precursor cluster (S-TGC precursor 2). Interestingly, there was a distinct high scoring cluster defined by expression of stem cell antigen-1 (*Sca1,* Sca1+ Trophoblast) (**Figure 4G-H**). In the literature, this population is described as a mid-gestational multipotent proliferative trophoblast population with stem cell-like capabilities to differentiate into trophoblast populations in both the labyrinth and the junctional zone^48,49^. This finding further supports the hypothesis that breast cancer cells co-opt a *PHLDA2*-driven developmental program, particularly of a trophoblast stem cell-like population, to promote metastasis.

### *PHLDA2* promotes vascular permeability

We next utilized immunofluorescence to investigate how *PHLDA2* overexpression may remodel tumor vasculature. Previous work has demonstrated that increases in tumor microvessel density correlates with an elevated risk of metastasis across various cancer types^47^. To assess vessel morphology and density, we performed CD31 immunofluorescent staining on *PHLDA2*-overexpressing and control HCI010 tumors. We observed no significant differences in vessel number, length, or microvessel density, suggesting *PHLDA2* may affect vascular function rather than structure (**Figure S5A-B**).

Our bulk RNA sequencing revealed that *PHLDA2* overexpression upregulated several genes within vascular development pathways known to regulate endothelial barrier integrity, including *SPARC*, which notably enhances melanoma lung metastasis by increasing vascular permeability^30^. Based on this, we hypothesized that PHLDA2 expression in tumor cells impairs endothelial barrier function. To test this functionally, we first employed an *in vitro* vascularized micro-tumor (VMT) assay, which recapitulates tumor–vessel interactions under physiologic flow^26–28^ (**Figure 5A-B**). VMTs were generated by embedding human fibroblasts, endothelial cells, and cancer cells (MDA-MB-231 GFP, *PHLDA2*-GFP, shScramble-GFP, or sh*PHLDA2*-GFP) in a fibrinogen matrix polymerized within microtissue chambers (**Figure 5A-B**). Culture medium was introduced through laminin-lined microfluidic channels to support vessel formation under interstitial flow. After four to five days, by which time a capillary network forms connecting the microfluidic “artery” and “vein”, 70 kDa fluorescent dextran was perfused through the microvessels, and permeability was quantified by fluorescence imaging of dextran leakage (**Figure 5A-B**). We found that *PHLDA2* overexpression in the tumor cells resulted in a two-fold increase in vessel permeability *in vitro*, while *PHLDA2* knockdown significantly reduced vessel permeability (**Figure 5C-D**).

**Figure 5:**
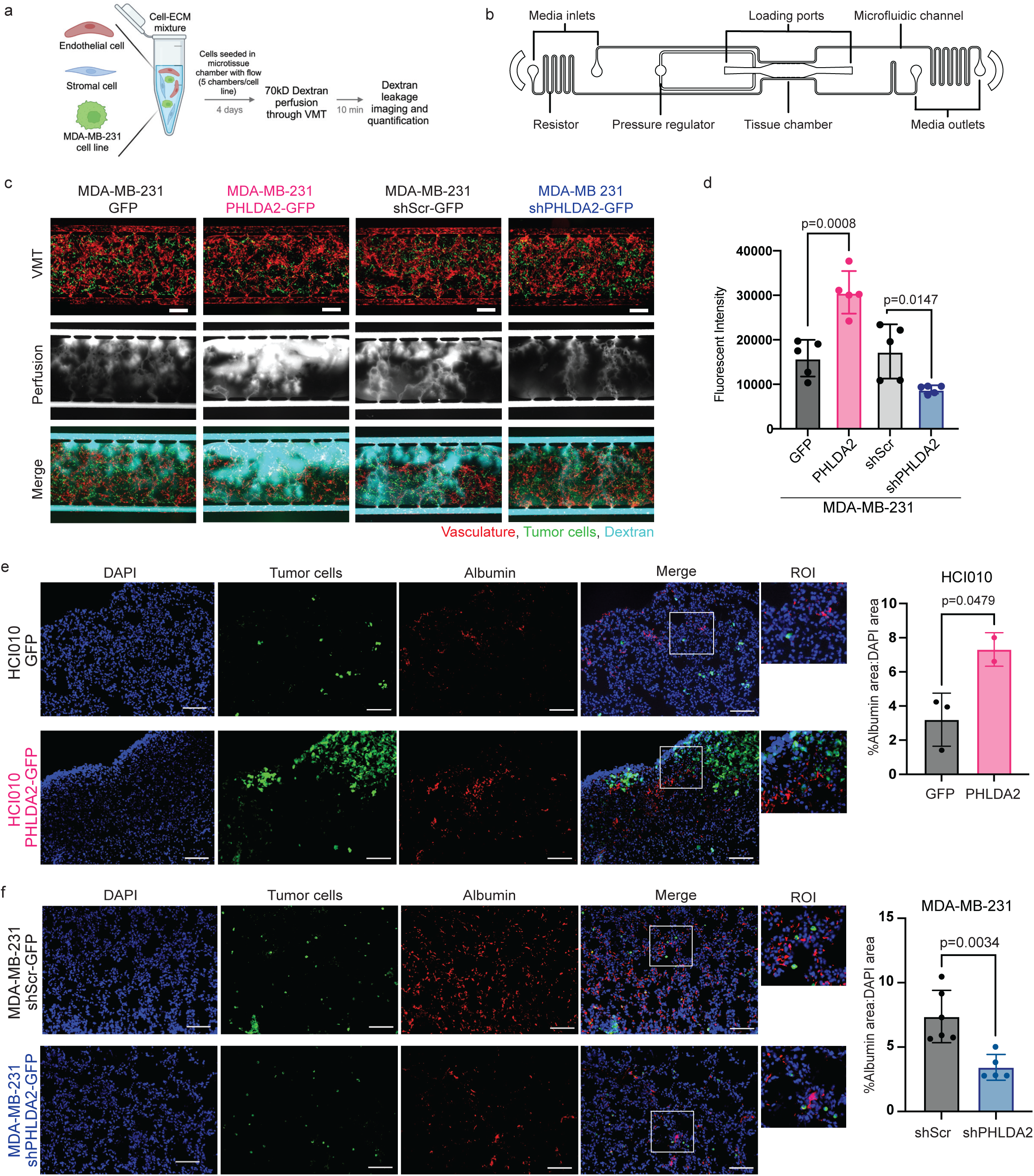
*PHLDA2* increases vascular permeability. (a) Schematic shows experimental design for evaluation of the effect of PHLDA2 expression on vascular permeability using the vascularized micro-tumor (VMT) model system. Human endothelial cells, stromal cells, and GFP-labeled MDA-MB-231 cancer cells were collected from 2D monolayer cultures and combined into a microtissue chamber with extracellular matrix to establish vascularized microtumors. After 4 days, 70kD fluorescence conjugated dextran was perfused through the VMT and imaged after 10 minutes for quantification. (b) Schematic illustrates the microfluidic device used for the VMT. The device includes a single 6 mm-long tissue chamber with medium inlets and outlets. The cell-ECM mixture was loaded into the tissue chamber through two ports. Loading is facilitated by a pressure regulator, and tissues are supported by hydrostatic pressure created across microfluidic channels connecting the media wells. Microfluidic resistors maintain physiological flow rates. (c) Representative fluorescent images display vascular leakage observed in each sample condition. MDA-MB-231 tumor cells (green) and vasculature (CD31, red) are shown in the top panels. Middle panels show vascular leakage following perfusion of 70 kD dextran (cyan) into the VMTs on day 4. The bottom panels show merged images. Scale bar = 500 μm. (d) Bar graph plotting quantification of dextran leakage for each sample condition in the VMT (n=5/group). Total extravascular fluorescence intensity (mean grey value) of dextran was quantified with ImageJ. Each point represents on VMT value obtained by the average of multiple 10x objective microscopic fields. Graph is displayed as mean ± SD. *P* values were calculated using multiple student’s t-tests. (e) Representative images of immunofluorescence staining for albumin (red) with DAPI (blue) and tumor cells (green) in spontaneous lung metastases from mice transplanted with HCI010 GFP and HCI010 *PHLDA2* tumor cells. Scale bar = 50 μm. Percent of albumin^+^ area normalized to DAPI^+^ area was quantified (n= 2-3/group). Each point represents one lung tissue value obtained by the average of 5-10 20x objective microscopic fields. Graph is displayed as mean ± SD. *P* values were calculated using student’s t-tests. (f) Representative images of immunofluorescence staining for albumin (red) with DAPI (blue) and tumor cells (green) in metastatic lung tissue from mice intravenously injected with MDA-MB 231 Scrambled GFP (control) or sh*PHLDA2* GFP. Scale bar = 50 μm. Percent of albumin^+^ area normalized to DAPI^+^ area was quantified (n= 5-6/group). Each point represents one lung tissue value obtained by the average of 5-10 20x objective microscopic fields. Graph is displayed as mean ± SD. *P* values were calculated using student’s t-tests.

To validate these findings *in vivo*, we performed albumin immunofluorescent staining on primary tumors and metastatic lungs to quantify vascular permeability. Albumin leakage into tissues serves as a surrogate marker for vessel permeability, as its large size prevents its passage through intact vessel barriers^50^. In HCI010 primary tumors, we found a marked increase in vascular permeability (**Figure S5C-D**). While in metastatic lungs from mice bearing HCI010 *PHLDA2*-overexpressing primary tumors, we observed a pronounced effect, with nearly 2.3-fold higher levels of albumin leakage (**Figure 5E**). Conversely, albumin leakage was reduced by approximately half in lungs containing MDA-MB-231 *PHLDA2* knockdown experimental metastases compared to controls (**Figure 5F**). Together, these data demonstrate that PHLDA2 expression in tumor cells robustly promotes vessel permeability in both *in vitro* and *in vivo* models, revealing a novel mechanism by which PHLDA2 may promote metastasis likely through enhancement of extravasation and intravasation.

### *SPARC* mediates *PHLDA2*-induced alterations in vascular permeability

Having established that *PHLDA2* promotes vascular permeability, we next investigated the underlying mechanism. We focused on the secreted glycoprotein SPARC as a potential mediator, given its established role in increasing vascular permeability and promoting metastasis in melanoma mouse models^30^. This hypothesis was strongly supported by our bulk RNA sequencing data, which identified *SPARC* as one of the most highly upregulated genes upon *PHLDA2* overexpression (log_2_ fold change = 4.44) (**Figure 6A**). We validated this link at both the transcript and protein levels. Quantitative PCR confirmed that *PHLDA2* overexpression significantly increased *SPARC* mRNA levels (**Figure 6B**). Consistently, Western blot analysis showed that *PHLDA2* overexpression markedly increased *SPARC* protein, while *PHLDA2* knockdown reduced its expression (**Figure 6C, S6A**). These results confirm that PHLDA2 robustly regulates SPARC. To determine if SPARC expression mediates PHLDA2-induced vascular permeability, we engineered MDA-MB-231 cells with *PHLDA2* knockdown vectors alone (sh*PHLDA2*-GFP) or in combination with *SPARC* overexpression (sh*PHLDA2*-GFP + *SPARC*-mCherry). In parallel, we generated a control cell line (shScramble-GFP mCherry) and a *SPARC* overexpression cell line (*SPARC*-mCherry). Successful knockdown and overexpression were confirmed at the protein level with Western blots (**Figure S6A**). We introduced these cell lines into the VMT system to assess changes in vascular permeability. Consistent with our previous experiments, knockdown of *PHLDA2* reduced vascular permeability (**Figure 6D-E**). However, the addition of *SPARC* overexpression increased vascular leakage back to levels observed in the control cell line (**Figure 6D-E**). These results demonstrate that SPARC mediates PHLDA2-driven vascular permeability and can restore permeability when PHLDA2 is suppressed.

**Figure 6:**
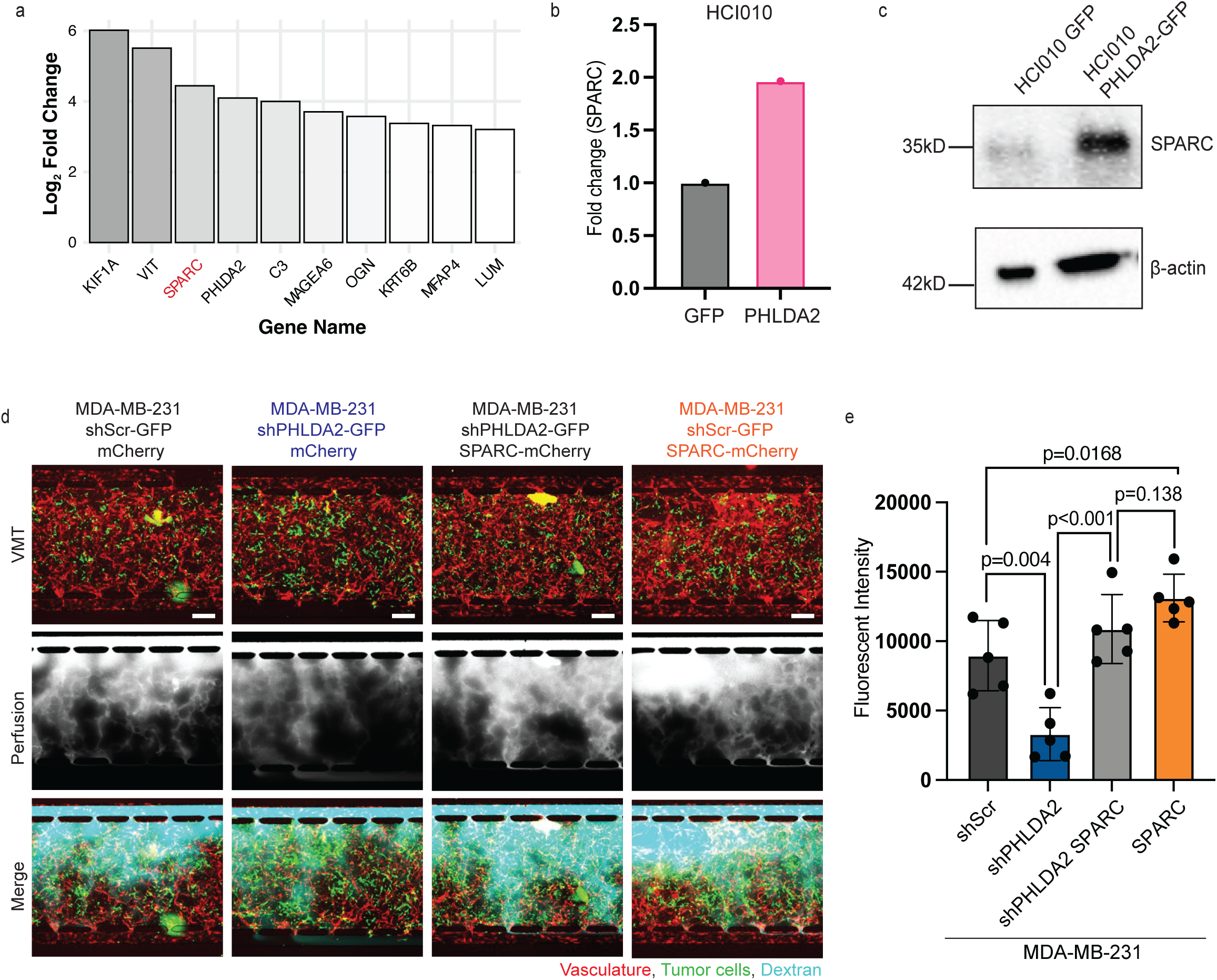
*SPARC* mediates *PHLDA2*-induced vascular leak. (a) Top ten upregulated genes in HCI010 *PHLDA2*-overexpressing tumor cells identified by bulk RNA sequencing. Bars represent the log_2_ fold change in gene expression compared to GFP control tumors. *SPARC* (highlighted in red) is among the significantly upregulated genes. (b) Bar graph plotting *SPARC* mRNA expression measured by quantitative PCR in HCI010 *PHLDA2* tumor cells (n=1) compared to HCI010 GFP controls (n=1). Gene expression is normalized to housekeeping gene control, GAPDH, and displayed as fold change relative to HCI010 GFP. (c) Western blot shows SPARC protein expression in HCI010 GFP (control) and *PHLDA2*-GFP cells that were sorted from primary tumors by flow cytometry. β-actin serves as a loading control. (d) Representative fluorescent images display vascular leakage observed in each sample condition. MDA-MB-231 tumor cells (green) and vasculature (CD31, red) are shown in the top panels. Middle panels show vascular leakage following perfusion of 70 kD dextran (cyan) into the VMTs on day 4. The bottom panels show merged images. Scale bar = 500 μm. (e) Bar graph plotting quantification of dextran leakage for each sample condition in the VMT (n=5/group). Total extravascular fluorescence intensity (mean grey value) of dextran was quantified with ImageJ. Each point represents on VMT value obtained by the average of multiple 10x objective microscopic fields. Graph is displayed as mean ± SD. *P* values were calculated using multiple student’s t-tests.

## Discussion

In this study, we identify the imprinted developmental gene *PHLDA2* as a potent driver of breast cancer metastasis. We provide evidence that *PHLDA2* is sufficient to promote lung metastasis in preclinical models and that its expression in patient tumors correlates with reduced metastasis-free survival. A central finding of our work is that *PHLDA2* achieves this through a transcriptional shift in breast cancer cells, making them more similar to a *Phlda2* expressing trophoblast population with high levels of *Sca1*. Based on literature, Sca1+ trophoblasts represent a stem-like population within the chorion and the labyrinth layer of the placenta mid-gestation (starting around E8.5)^48^. Their stem-like nature is defined by their ability to self-renew and their multipotency, as they can be isolated and differentiated into various trophoblast subtypes of both the labyrinth and junctional zone. Additionally, trophoblasts stem cells co-expressing CD117 and Sca1 have robust multipotency with the ability to differentiate into lung alveolar epithelial cells, cardiomyocytes, and retinal photoreceptor cell^51^. This is particularly compelling parallel as it is well-established that the cancer progression demands cellular plasticity^4^. Furthermore, our findings suggest a mechanism for the aberrant reactivation of *PHLDA2*. We observed an inverse correlation between *PHLDA2* expression and methylation at specific exonic CpG sites indicates that epigenetic dysregulation, likely a loss of imprinting, unlocks this latent developmental program in adult cancer cells^1,3^. Therefore, we speculate that *PHLDA2* overexpression may confer a trophoblast stem cell-like state, reactivating a developmental program that unlocks the phenotypic plasticity that cancer cells required to successfully navigate the challenges of metastasis.

Our study additionally elucidates pro-metastatic functions downstream of *PHLDA2* including ECM and vascular remodeling. Specifically, *PHLDA2* overexpression increases collagen and MFAP5 deposition in metastatic lesions and increases vascular permeability in primary tumors and lungs. We further identified the secreted matricellular protein, SPARC, as a key mediator of this vascular effect, consistent with previous work on SPARC in other cancers^30^. The impact of *PHLDA2* on vascular remodeling is again reminiscent of placental trophoblasts. During normal placental development, invasive trophoblasts are responsible for profound vascular remodeling. They invade the maternal spiral arteries, replacing the endothelial and smooth muscle lining^52^, and promote angiogenesis within the labyrinth to create the vascular network essential for fetal nourishment^14^. This process requires a precise balance of gene expression to ensure proper vessel integrity and function. Our findings suggest that the level of *PHLDA2* is critical to this balance. While normal *PHLDA2* levels in trophoblasts likely contribute to controlled vascular remodeling, the aberrant overexpression we modeled in cancer cells may drive excessive vascular permeability. This speculation aligns with clinical observations in humans, where high placental *PHLDA2* expression is strongly correlated with preelampsia^53^ and fetal growth restriction^12^, conditions linked to abnormal vascular development^54–56^. Therefore, the increased vascular permeability we observe in tumors may represent a pathological manifestation of a developmental process that, when dysregulated in the placenta, leads to compromised nutrient exchange and poor fetal outcomes.

While our study sheds new light on the roles of PHLDA2 in breast cancer, certain limitations offer avenues for future investigation. The precise molecular mechanism linking PHLDA2, a protein with a PH domain, to the transcriptional upregulation of genes like *SPARC* remains unknown. PH domains interact with phosphoinositides (PIPs) in the plasma membrane and can function in potentiating signal transduction or act as scaffolds^57^. Our bulk sequencing revealed hundreds of upregulated genes in response to *PHLDA2* overexpression; thus, it is plausible that PHLDA2 interacts with lipids on the cell membrane to promote signal transduction resulting in the transcription of a pro-oncogenic meta-program. Future studies employing techniques such as Co-Immunoprecipitation would identify proteins in which PHLDA2 interacts with and reveal how it drives this gene program. Additionally, exploring the potential therapeutic efficacy of targeting PHLDA2 axis including potential upstream epigenetic factors which reduce methylation at *PHLDA2* gene locus in breast cancers or downstream pro-metastatic regulators including SPARC is a critical next step.

In conclusion, our study establishes PHLDA2 as a critical driver of breast cancer metastasis by hijacking a developmental program reminiscent of placental trophoblasts. By promoting ECM remodeling and SPARC-mediated vascular permeability, PHLDA2 creates a microenvironment highly permissive for metastatic dissemination. These findings not only provide profound insight into the fundamental mechanisms of metastasis but also highlight PHLDA2 as a promising prognostic biomarker and a potential therapeutic target. Ultimately, disrupting the aberrant reactivation of such developmental pathways may offer a novel strategy to combat metastatic disease.

## Materials and Methods

### Mouse placenta spatial transcriptomics data analysis

Mouse placenta spatial transcriptomics data previously published^32^ was analyzed for PHLDA2 expression using the interactive tool provided by the authors (https://db.cngb.org/stomics/mpsta/). Cell annotations assigned in original study were used.

### Mouse placenta single cell RNA sequencing data analysis

The single-cell RNA sequencing dataset, along with its corresponding metadata, was downloaded from the Gene Expression Omnibus (GEO) under the accession number GSE156125^35^. File was loaded into Seurat v4.3.0 in R version 4.2.2. For the initial phase of our analysis, we utilized the original cell embeddings and annotations provided in the publication associated with this dataset^35^. We subset the clustered identified as trophoblasts, parietal trophoblast giant cells (P-TGCs), and spiral artery trophoblast giant cells (SpA-TGCs) for more detailed investigation. This trophoblast-specific subset was then subjected to a new round of clustering to achieve a higher resolution of trophoblasts cell states. Following re-clustering, the cell types were re-annotated. This re-annotation was performed manually by examining the expression of established trophoblast subtype marker genes, which were extensively described in the original publication^35^. Markers included: *Eomes* and *Lin28a* for trophoblast stem cells (TSC) and the extraembryonic ectoderm (ExE), *Met* for labyrinth trophoblast progenitor cells (LaTP), *Gjb5*, *Slc16a3*, and *Mt2* for ectoplacental cone cells (EPC), *Cts7*, *Adamts1*, and *Fmnl1* for EPC migratory cells, *Prl7b1* and *Pcdh12* for glycogen trophoblasts (GlyT), *Prl3b1*, *Prl7a2*, *Prl8a8* for spiral artery trophoblasts giant trophoblast cells (SpA-TGC), *Ugcg* and *Prl3d1*^low^ for primary parietal trophoblast giant cells (primary P-TGC), *Ugcg* and *Prl3d1*^high^ for secondary parietal trophoblast giant cells (secondary P-TGC), *Krt19*, *Timp1*, and *Phactr1* for sinusoidal trophoblast giant cells precursors (S-TGC precursor), *Tfrc* for syncytiotrophoblast I precursors (SynTI precursor), *Gcm1* for syncytiotrophoblast II precursors (SynTII precursor), Ctsq^high^ for sinusoidal trophoblast giant cells (S-TGC), and *Ascl2*^high^ for spongiotrophoblasts (SpT).

Gene scoring analysis was performed on *Phlda2*-expressing trophoblasts which were subset based on the threshold of *Phlda2* expression greater than 0.1. After subset, 76% of trophoblasts remained demonstrating the majority of trophoblasts across placental development have detectable *Phlda2* expression. We generated a *PHLDA2*-overexpressing tumor cell gene signature comprising of the top 100 upregulated differentially expressed genes from our bulk RNA sequencing (**Table 2**). The signature was converted to mouse genes using the R package babelgene (v22.9) and scored in the subset *Phlda2*-expressing trophoblasts using the AddModuleScore function in Seurat.

### Relapse-Free Survival Analysis

To assess relapse-free and distal metastasis-free survival, we used the KM plotter database to generate Kaplan-Meier survival curves using a primary tumor microarray dataset of breast cancer patients. All Kaplan-Meier plots use ‘auto select best cutoff’ parameter.

### TCGA Gene Expression Analysis

To assess gene expression in primary tumor versus normal tissue, the UALCAN analysis portal was used to mine the TCGA Breast Invasive Carcinoma dataset. Statistics were provided by the portal.

### Analysis of epigenetic dysregulation of PHLDA2 in breast cancer

Bulk methylation and gene expression profiles for the PHLDA2 gene were obtained from the GDC TCGA Breast Cancer repository. Exploratory data analysis and data download was performed with the UCSC Xena platform^58^. In-depth analysis was performed on the singularity container server “Bioportal1” with a JupyterLab instance created by the UC Irvine Research Cyberinfrastructure Center. An R notebook with the following packages was used: R (v4.1.2), ggplot2 (v3.3.5), dplyr (v.1.0.8), reshape (1.4.4). Primary tumors and normal samples which contained missing data were filtered out using “na.omit”. For single-modality plotting, only data for that modality was considered in the filtration. For joint methylation and gene expression analysis, samples with missing data in either modality were excluded.

To select the candidate regulatory sites for PHLDA2, the methylation beta values of all 32 associated loci from Infinium HumanMethylation450 BeadChip were visualized as box and whisker plots. Welch’s t-test from UCSC-Xena as well as visual examination of the differences in the means in the tumor and normal samples for each locus identified two loci as most differentially demethylated in breast cancer (cg05167973 and cg04720330). These loci were replotted with their adjusted p-values after a Bonferroni multiple testing adjustment and retained for further analysis.

To test whether the two demethylated loci have a role in upregulating *PHLDA2* expression in breast cancer, the methylation beta values of each locus was plotted against the RNA expression of *PHLDA2* on a scatter plot. The negative sign of the slope for the best-fit line as well as Spearman’s rank correlation rho were used to infer the inverse relationship between methylation levels at the candidate regulatory loci and the levels of gene expression. T-tests and Bonferroni p-value adjustments were performed again.

### PDX Sample Propagation

A patient derived xenograft (PDX) sample was provided by A. L. Welm at the Department of Oncological Sciences at the Huntsman Cancer Institute (HCI). Tissues were collected from individuals being treated at the Huntsman Cancer Hospital and University of Utah with informed consent under a protocol approved by the Institutional Review Board of the University of Utah. A pleural effusion of a female patient diagnosed with Stage IIIC ER−PR−Her2− basal-like (PAM50) invasive ductal carcinoma and treated with several rounds of chemotherapies is noted as HCI-010. This sample was collected and de-identified by the Huntsman Cancer Institute Tissue Resource and Application Core facility before being obtained for implantation. The study is compliant with all of the relevant ethical regulations regarding research involving human participants.

### Cell Culture

Breast cancer cell line MDA-MB-231, was cultured in DMEM, 1X (Corning, 10-013-CV) media respectively with 10% heat-inactivated fetal bovine serum (Sigma-Aldrich, cat. no. 12306 C) and 1% penicillin/streptomycin (Hyclone, SC30010) 100 X solution. The cells were passaged with 0.05% trypsin (Corning, 25-052-Cl) after reaching 70% confluency. MDA-MB-231 cells are mycoplasma-free and STR profiled with a 93% match to the ATCC reference.

### Viral Transduction (PDX and Cell Lines)

PDX cells were transduced as previously described^41^. GFP control (+GFP) and human PHLDA2-GFP (+*PHLDA2*-GFP) lentiviral expression vectors purchased from VectorBuilder Inc., Chicago, IL, USA were packaged into lentiviral particles. MDA-MB-231 cell line was transduced with Scramble GFP control (+Scramble-GFP) and human PHLDA2 shRNA-GFP (+sh*PHLDA2*-GFP) lentiviral expression vectors purchased from VectorBuilder Inc., Chicago, IL, USA were packaged into lentiviral particles. All cell lines were transduced with 10 mg/mL polybrene. Cells were incubated in virus overnight before switching to fresh media to allow cells to grow out. GFP positive transduced cells were selected using FACS. Alterations in PHLDA2 expression were confirmed by qPCR and Western Blot Analysis.

### Animal Experiments

For spontaneous metastasis models, HCI010 (120,000) cells were mixed 1:1 with sterile PBS and Matrigel (Corning 356230) and injected orthotopically into the number four mammary fat pad area of female NSG mice using a 0.5 mL insulin syringe. For experimental metastasis models, cultured MDA-MB-231 (500,000 cells) cells were resuspended in 100 uL of sterile PBS and injected intravenously into the tail vein of female NSG mice. Primary tumors were measured using a caliper and volumes were calculated using the tumor volume = 1/2 (length x width^2^) equation. All animal experiments were reviewed and approved by The Institutional Animal Care and Use Committee of the University of California, Irvine.

### Tissue Harvest and Dissociation

At the endpoint, animals were euthanized by asphyxiation with CO_2_ followed by cervical dislocation and perfusion with 10 mM EDTA in PBS. Tumors and lungs were removed and dissociated for flow cytometry by mechanical chopping with razor blades. The chopped tissues were digested in 2 ug/mL Collagenase IV (Sigma-Aldrich cat. No. C5138-1G) in DMEM with 300 mg/mL DNAse I (Catalog Number) for 20 minutes at 37°C. The cell suspensions were washed with Hanks Balanced Salt Solution (HBSS) and passed through a 70 μm cell strainer. Lung and primary tumor cells were treated with 1X Red Blood Cell Lysis buffer followed by resuspension in PBS for FACS analysis.

### Flow Cytometry

Cells were stained with a fluorescently conjugated antibodies for CD298/MHCI to isolate human cells. Human specific-CD298 (diluted 1:100; PE; BioLegend, cat. no. 341704) and the mouse-specific antibody MHC-I (diluted 1:150; APC; Thermo Fisher Scientific, cat. no. 17-5957-80) antibodies were added to samples. Cell viability was determined using SYTOX blue stain (diluted 1:1000, Thermo Fisher Scientific, cat. No. S34857). Cell viability was determined using SYTOX blue stain (diluted 1:1000, Thermo Fisher Scientific, cat. no. S34857). The samples were resuspended in PBS for FACS analysis using the BD FACS Aria Fusion cell sorter (Becton, Dickinson and Company, Franklin Lakes, NJ, USA). To isolate single cells from doublet and multiplet cells, forward scatter area by forward scatter width (FSC-A x FSC-H) and side scatter area by side scatter width (SSC-A x SSC-H) were used. GFP labeled human primary tumor and metastatic lung cells were selected by gating on Sytox^-^GFP^+^ cells.

### Quantitative PCR

Prior to RNA extraction, HCI010 tumors were dissociated into a single cell suspension and sorted to isolate GFP^+^ tumor cells or MDA-MB-231 cells were trypsinized from a cell culture dish. RNA was extracted from cells using Quick-RNA MicroPrep (Zymo Research, R1050). cDNA was obtained from iScript cDNA Synthesis Kit (Bio Rad, 1708891). qPCR amplification was performed using Power SYBR Green PCR Master Mix (Applied Biosystems, A25742). Gene specific primers were utilized to amplify *PHLDA2* (Forward Primer: CCGCCGCGGGCCATA, Reverse Primer: CCACAGCCGGATGGTAGAAA), *MFAP5* (Forward Primer: AGTCAACGAGGAGACGATGTG, Reverse Primer: CATCCCAGCACTCCAAGTCA), S*PARC* (Forward Primer: TGAGAATGAGAAGCGCCTGG, Reverse Primer: TGGGAGAGGTACCCGTCAAT), *COL1A1* (Forward Primer: ACAAGGCATTCGTGGCGATA, Reverse Primer: ACCATGGTGACCAGCGATAC), and *LUM* (Forward Primer: CCGTCCTGACAGAGTTCACA, Reverse Primer: TGGCAAATGGTTTGAATCCTTACT). Each sample was normalized to the housekeeping gene *GAPDH* (Forward Primer: CTCTCTGCTCCTCCTGTTCGACGAC, Reverse Primer: TGAGCGATGTGGCTCGGCT). Primers were generated by Integrated DNA Technologies. Relative quantification of transcripts for all samples was performed in triplicates.

### Western Blot

Prior to cell lysis, HCI010 tumors were dissociated into a single cell suspension and sorted to isolate GFP^+^ tumor cells or MDA-MB-231 cells were trypsinized from a cell culture dish. Cells were lysed in RIPA buffer (150mM NaCl, 50mM Tris-HCl pH 8.0, 1% Triton-X, 0.5% sodium deoxycholate, 0.1% SDS, diluted in H_2_O) containing protease and phosphatase inhibitors (Thermo Fisher, Cat No. 78430). Lysates were incubated with agitation for 45 minutes and centrifuged at 12,000g for 10 minutes. Protein lysate supernatant was removed and quantified using Pierce BCA Protein Assay Kit (Thermo Fisher, Cat No. 23225) according to manufacturer’s instructions. 8-20ug protein samples were run on a 12% SDS PAGE gel (BioRad, Cat No. 4568046) and transferred to a PVDF membrane using a wet transfer protocol. Primary antibodies were diluted in a blocking solution of 5% BSA in TBS containing 0.1% Tween-20 (5% BSA-TBST) and incubated at 4°C overnight. Primary antibodies included: 1:500 Rabbit anti-Human TSCC3 (PHLDA2) antibody (Abcam, Cat No. ab234669), 1:1000 Rabbit anti-Human SPARC antibody (Thermo Fisher, Cat No. PA5-78178), 1:1000 Mouse anti-Human β-actin (Santa Cruz, Cat No. sc-47778), and 1:1000 Mouse anti-Human Vinculin (Millipore Sigma, Cat No. V9131). Secondary antibodies were diluted in blocking solution and placed on membranes for one hour at room temperature. Secondary antibodies included: 1:1000 Goat Anti-Mouse HRP (Thermo Fisher, Cat No. 31430) or 1:1000 Goat Anti-Rabbit HRP (Thermo Fisher, Cat No. 31460). Protein bands were visualized using Thermo Fisher Chemiluminescent Substrate kit (Thermo Fisher, Cat No. 34579) and imaged on a BioRad ChemiDoc system. Densitometry was performed using ImageJ and protein expression was normalized to β-actin or Vimentin.

### Tissue Preparation

Tumor and lung tissues for Masson’s trichrome staining were fixed in 10% neutral-buffered formaldehyde at room temperature for 24 hours, placed in freshly prepared 70% ethanol, and process for paraffin embedding in a Leica tissue processor using standard protocols. FFPE blocks were sectioned into 5 μm sections using a Reichert-Jung 2040 Autocut Rotary Microtome and stained with Masson’s trichrome protocol.

Tumor and lung issues for immunofluorescence staining were fixed in formaldehyde for 6-12 hours at 4℃ and placed in 30% sucrose overnight. Tissues were embedded with OCT Compound (Tissue-Tek, Cat no. 4583) in disposable base mold at -80℃. Tissue sections of 8 μm were sectioned using a Leica Cryostat CM1950 and stored in a closed container at -80℃.

### Immunofluorescence staining

Slides with OCT-embedded tissue sections were washed, blocked with BlockAid Blocking Solution (Life Technologies Corporation, Cat no. 810710) at room temperature for an hour, and incubated overnight with primary antibodies (1:100 Ki67, Genetex, Cat No. GTX16667, 1:100 MFAP5 Rabbit Polyclonal Antibody, Proteintech, Cat no. 15727-1-AP, CD31 Rat Anti-mouse, 1:75, BD Bioscience, Cat no. 741740) at 4℃. Albumin Rabbit Polyclonal Antibody (1:100, Proteintech, Cat no. 16475-1-AP) was used in the same method but blocking occurred in 5% Fish Gelatin. Tissues were incubated with appropriate secondary antibodies for 1 hour in the dark at room temperature. Tissues were mounted with Antifade Mounting Medium with DAPI (Epredia, Cat no. 8312-4) and sealed with a coverslip. Slides were imaged on Keyence BZ-X series inverted microscope.

### Immunofluorescence Image Quantification

Five-ten 20x regions of interest were captured per tissue stained. Each image was quantified and all images corresponding to a sample were averaged to report one value per mouse on graphs. The following analyses were performed using NIH open-source image software ImageJ (http://rsbweb.nih.gov/ij/).

### Ki67^+^ GFP^+^ area

Ki67^+^ GFP^+^ double positive area was quantified by first using thresholding to measure the area of GFP^+^. GFP^+^ area was converted to a mask and projected onto the same image’s Ki67^+^ channel. Ki67^+^ area across the image was visualized using a constant threshold. Next, Ki67^+^ area within the GFP^+^ mask was measured. Lastly, Ki67^+^ GFP^+^ area was normalized to total GFP^+^ area and multiplied by 100 to measure the percent of Ki67^+^ area within GFP^+^ metastases or the Ki67^+^ GFP^+^ area. We selected this method specifically to quantify Ki67-positive tumor cells, while avoiding the inclusion of other proliferating cell types in the metastatic lung.

### MFAP5 expression and Albumin leakage

MFAP5 and albumin staining were quantified as the ratio of MFAP^+^ or albumin^+^ area normalized to DAPI^+^ area and multiplied by 100 to quantify percent of MFAP or albumin area relative to DAPI area. We determined MFAP5, albumin, or DAPI area using consistent threshold values across samples. We chose this method of quantification to determine the expression of MFAP5 or leakage of albumin relative to total lung tissue area.

### Vessel Structure Analysis

Tumor vessel structural analysis was performed as described previously^59^. Briefly, vessel parameters quantified included the total number of CD31^+^ vessels, the average vessel length of all vessels (reported in μm), the number of vessels with a length greater than 50 μm (elongated vessels), the number of open lumens, and the number of 100 μm^2^ regions per image occupied by CD31^+^ vessels (microvessel density).

### Masson’s Trichrome Staining and Quantification

Tissues were deparaffinized with Histo-Clear and decreasing concentrations of ethanol (100%, 95%, 70%) for one minute each. Each step was done twice. Masson’s trichrome staining was performed using Trichrome Stain Kit (Abcam, Cat no. ab150686) according to manufacturer instructions. Briefly, Bouin’s fluid was added onto tissues and incubated for one hour, followed by a wash step. Equal parts of Weigert’s reagents A and B were mixed to create Weigert’s Iron Hematoxylin and added onto tissues for five minutes and then washed. Tissues were then incubated with a Biebrich Scarlet/Acid Fuchsin solution for 15 minutes followed by a wash step. After, tissues were incubated in Phosphomolybdic/Phosphotungstic Acid solution for 15 minutes. Without washing Aniline Blue was immediately added to the slides and incubated for 10 minutes. Slides were then washed and incubated in a 1% Acetic Acid solution for 5 minutes. Tissues were then dipped in 95% ethanol and 100% ethanol and left in Histo-Clear (National Diagnostics, HS-200) for one minute. Tissues were mounted with a mounting medium (Cytoseal XYL, Epredia, 8212-4) and coverslip. 10x objective images were acquired on the Keyence BZ-X series inverted microscope. Open-access image software, QuPath (https://qupath.github.io/), was used for analysis. In QuPath, a border was drawn around a metastatic lesion to create a region of interest (ROI) and area of the ROI was measured. Blue pixels (collagen) were then isolated from other colors using the QuPath’s color deconvolution stains and the area of collagen within the metastatic lesion ROI was measured. Lastly, the area of collagen within a metastatic lesion was normalized to total area of metastatic lesion and multiplied by 100 to quantify the percent of collagen deposition in a metastatic lesion. This process was repeated for every lesion identified in 5-10 randomly acquired 10x objective brightfield images. Each lesion was plotted as a point on the graph. In total, 3 lungs were imaged per group.

### Cell Growth Assays (Cell Titer-Glo 3D Cell Viability Assay, MTT Cell Proliferation Assay)

HCI010 PDX tumor organoids were cultured in EpiCult-B Basal Medium (Mouse, StemCell, 05611) supplemented with 10 ng/mL FGF and EGF recombinant protein. After 7 days, HCI010 PDX tumor organoids were manually broken apart with pipetting. 200,000, 500,000 or 100,000 cells were seeded in a 24 well ultra-low attachment plate to allow spheroids to grow. Cell Titer-Glo 3D Cell Viability Assay (Promega, Cat no. G9681) protocol was followed as directed by the manufacturer and imaged using the luminescence function on Biotek Cytation 5 plate reader.

MDA-MB-231 cells were seeded at 5,000 cells/well in a standard flat bottom 96 well plate. MTT Cell Proliferation Assay Kit (Cayman Chemical, Cat no. 10009365) protocol was followed as directed by manufacturer and imaged using absorbance function on Biotek Cytation 5 plate reader.

### Bulk RNA-Sequencing Library Prep

RNA was extracted from sorted primary tumor cells (Viable GFP^+^) using Quick-RNA MicroPrep (Zymo Research, R1050) as directed by the manufacturer. Total RNA was monitored for quality control using the Agilent Bioanalyzer Nano RNA chip and Nanodrop absorbance ratios for 260/280nm and 260/230nm. Library construction was performed according to the Illumina TruSeq mRNA stranded protocol. The input quantity for total RNA within the recommended range and mRNA was enriched using oligo dT magnetic beads. The enriched mRNA was chemically fragmented. First strand synthesis used random primers and reverse transcriptase to make cDNA. After second strand synthesis the ds cDNA was cleaned using AMPure XP beads and the cDNA was end repaired and then the 3’ ends were adenylated. Illumina barcoded adapters were ligated on the ends and the adapter ligated fragments were enriched by nine cycles of PCR. The resulting libraries were validated by qPCR and sized by Agilent Bioanalyzer DNA high sensitivity chip. The concentrations for the libraries were normalized and then multiplexed together. The multiplexed libraries were sequenced using paired end 100 cycles chemistry for the Illumina NovaSeq 6000. 50 million reads were sequenced per sample.

### Bulk Sequencing Analysis

Bulk RNA sequencing data was processed following a standardized workflow. Initial quality control of raw sequencing reads was conducted using FastQC, followed by alignment to the Homo sapiens reference genome (hg38) using HISAT2. Gene-level read counts were generated using featureCounts, counting only exonic regions. Differential gene expression analysis was performed with DESeq2, identifying differentially expressed genes (DEGs) with an FDR-corrected adjusted p-value < 0.05 and log_2_ fold change ≥ 0.5 for the *PHLDA2* overexpression samples compared to GFP control samples. Low-count genes were filtered out before calculating reads per kilobase per million mapped reads (RPKM). For further analysis, only protein-coding genes with an average reads per kilobase per million mapped reads (RPKM) above 2 were included in volcano plots. Volcano plot was plotted with R package EnhancedVolcano (v1.20.0). The gene expression heatmap displays normalized gene expression levels in transcripts per million (TPM). Heatmap was made using pheatmap (v1.0.12). Gene set enrichment analysis (GSEA) was performed using R packages clusterProfiler (v4.10.1) and org.Hs.eg.db (v3.18.0). Top ten significant (adjusted p value > 0.05) pathways of interest were selected based on biological relevance. Normalized enrichment score bar plots were generated with ggplot2 (v3.5.1). Enrichment plots were generated with enrichplot (v1.22.0).

### Microfluidic Device Fabrication for VMT

Microfluidic device fabrication and loading has been previously described^26–28^. In summary, a custom polyurethane master mold is created using a 2-part polyurethane liquid plastic (Smooth Cast 310, Smooth-On Inc.). Subsequently, a PDMS layer is replicated from this master mold, and holes are punched to create inlets and outlets. The platform is assembled in two stages: first, the PDMS layer is chemically glued and subjected to 2 minutes of oxygen plasma treatment to affix it to the bottom of a 96-well plate. Following this, a 150 µm thin transparent membrane is bonded to the bottom of the PDMS device layer through an additional 2-minute treatment with oxygen plasma. The fully assembled platform is then placed in a 60°C oven overnight, covered with a standard 96-well plate polystyrene lid, and sterilized using UV light for 30 minutes before cell loading.

### Cell Culture and Microfluidic Device Loading for VMT

To establish the vascularized microtumor (VMT), healthy human lung fibroblasts and ECFC-ECs (endothelial colony forming cell endothelial cells) were harvested and resuspended in fibrinogen solution at a concentration of 3×10^6^ cells/mL and 7×10^6^ cells/mL, respectively. MDA-MB-231 cells (GFP, *PHLDA2*, Scramble or sh*PHLDA2*) were introduced into the mixture at a concentration of 1 × 10^5^ cells/mL fibrinogen solution. Fibrinogen solution was prepared by dissolving 70% clottable bovine fibrinogen (Sigma-Aldrich) in EBM2 basal media (Lonza) to a final concentration of 5 mg/mL. The cell-matrix suspension (6 μL) was mixed with thrombin (50 U/mL, Sigma-Aldrich) at a concentration of 3U/mL, quickly seeded into each microtissue chamber, and allowed to polymerize in a 37◦C incubator for 15 minutes. Laminin (1 mg/mL, LifeTechnologies) was then introduced into the microfluidic channels through medium inlets and incubated at 37◦C for an additional 15 minutes. After incubation, culture medium (EGM-2, Lonza) was introduced into the microfluidic channels and medium wells. Medium was changed every other day and hydrostatic pressure head re-established daily to maintain interstitial flow.

### VMT Imaging and Analysis

Fluorescence images were acquired with a Biotek Lionheart fluorescent inverted microscope using automated acquisition and standard 10x air objective. To test vessel perfusion, 25 µg/mL rhodamine-conjugated 70 kDa dextran was added to the medium inlet for 10 minutes. ImageJ software was utilized to determine the total extravascular fluorescence intensity (mean grey value) for each VMT. Subtraction of background fluorescence measurements was performed for each chamber.

### Statistics and Reproducibility

Data are presented as the mean ± s.d. from two or three independent experiments, unless stated otherwise. Statistics were conducted using Student’s *t* test. Statistical test results are represented on graphs. No statistical method was used to determine sample size. Age-matched female NSG mice were breed in-house and randomly assigned into experimental groups. Investigators were aware which conditions applied to each experimental group while performing experiments and analyzing data.

## Acknowledgements

P.H. was supported by the UCI Achievement Rewards for College Scientists (ARCS) fellowship. H.S. was supported by the National Cancer Institute (NCI) under Award Number T32CA009054. S.J.H and C.C.W.H. were supported by the NIH U54 CA217278. T.L. was supported by the NCI under Award Number F31CA281331. R.D. was supported by the National Institute of Health (NIH) CSUF/UCI-CFCCC Cancer Health Disparities Research Program (P20CA253251 and P20CA25325). J.I-R. was supported by a Feodor Lynen Research Fellowship from the Alexander von Humboldt Stiftung. T.P.M was supported by the UCI Medical Scientist Training Program (NIH/NIGMS 1T32GM154637-01) and the CIRM Training Grant (CIRM EDUC4-12822). I.A. was supported by the UCI Medical Scientist Training Program (NIH/NIGMS 1T32GM154637-01), and the NCI under Award Number 5T32CA009054-44 and Award Number 1F30CA306157-01. K.K. was supported by the funding from the NIH/NCI (R01CA234496), NIH/NIGMS (R01GM147741) and by an Emerging Leader Award from the Mark Foundation for Cancer Research. D.M. was supported by a Canadian Institutes of Health Research Postdoctoral Fellowship award. D.M. and this work were supported by a NIH/NCI K99/R00 Pathway to Independence Award (1K99CA267160-01). This work was supported by an American Cancer Society (ACS) Research Scholar Grant (RSG) (134389-RSG-039-01-DDC), NIH/NCI grants (R01CA237376-01A1 and 1P01CA288662-01A1) awarded to D.A.L. Additionally, this work was supported by funding from Team Michelle. This work was also supported in part by an Opportunity Award (U54CA217378) funded by the NCI to the University of California, Irvine, Center for Complex Biological Systems (CCBS; NCI, grant no. U54CA217378) to D.M, T.L., K.K., and D.A.L.. This work utilized the infrastructure for high-performance and high-throughput computing, research data storage and analysis, and scientific software tool integration built, operated, and updated by the Research Cyberinfrastructure Center (RCIC) at the University of California, Irvine (UCI). The RCIC provides cluster-based systems, application software, and scalable storage to directly support the UCI research community (https://rcic.uci.edu). Methylation analysis was conducted with valued assistance by Jie “Jenny” Wu from the UC Irvine Department of Biological Chemistry and the UCI Genomics High Throughput Facility. We want to thank Shannon Pfeiffer for assisting with downloading TCGA data. We would like to acknowledge Pauline Nguyen and Vanessa Scarfone for all their help in the UCI Flow Cytometry Core. Additional thanks to Melanie Oakes and the UCI Genomic Core staff for assistance with the bulk RNA sequencing. Schematics were made using Biorender.

## Author Contributions

D.M. and D.A.L. conceptualized the study. The investigation was conducted by P.V.H., H.S., S.J.H., T.L., A.L.P.A., C.C., S.J., R.M.D., I.L., S.M., J.I-R., I.A., H.A., G.A.H., A.L., T.P.M., and D.M. The methodology was developed by P.V.H., H.S., S.J.H., T.L., I.L., A.L.P.A., C.C., S.J., R.M.D., and D.M. Data curation was performed by P.V.H., H.S., S.J.H., T.L., S.J., I.L., A.L.P.A., C.C., S.J., R.M.D., and D.M., while formal analysis was carried out by P.V.H., H.S., S.J.H., T.L., I.L., S.M., and D.M. The original manuscript was drafted by P.V.H., H.S., S.J.H., C.C., R.M.D., and D.A.L. All authors contributed to the review and editing of the manuscript. Supervision of the project was provided by C.C.W.G., K.K., D.M., and D.A.L. Project administration was managed by S.M. and J.G. Resources were provided by C.C.W.G., K.K., D.M., and D.A.L., and funding was acquired by T.L., D.M., K.K., and D.A.L.

## Data availability statement

Mouse placenta single cell RNA sequencing dataset was previously published^35^ and deposited on NCBI’s Gene Expression Omnibus at accession number GSE156125. Mouse placenta spatial transcriptomics dataset was previously published^32^ with an interactive data portal (https://db.cngb.org/stomics/mpsta/) which we utilized for this study. Data generated during this study are available from the corresponding author upon request. Our bulk RNA sequencing dataset is publicly available in NCBI’s Gene Expression Omnibus at accession number GSE279530.

