## Supplementary Figures for "PHLDA2 promotes breast cancer metastasis by co-opting a developmental program for placental vascular remodeling"

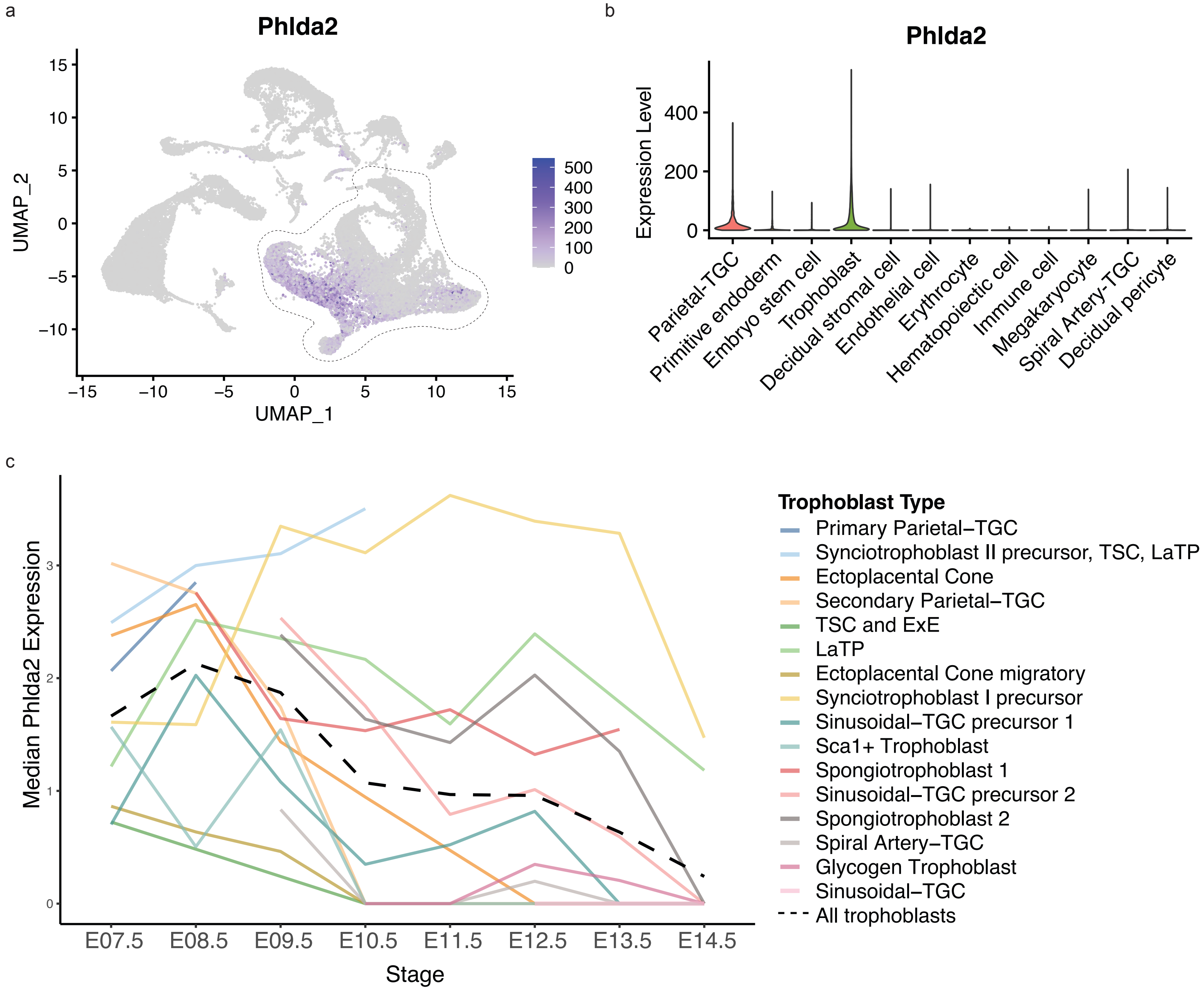
a
Phlda2
UMAP_2
UMAP_1
b
Phlda2
Expression Level
Parietal-TGC
Primitive endoderm
Embryo stem cell
Trophoblast
Decidual stromal cell
Endothelial cell
Erythrocyte
Hematopoietic cell
Immune cell
Megakaryocyte
Spiral Artery-TGC
Decidual pericyte
c
Median Phlda2 Expression
Stage
Trophoblast Type
Primary Parietal–TGC
Synciotrophoblast II precursor, TSC, LaTP
Ectoplacental Cone
Secondary Parietal–TGC
TSC and ExE
LaTP
Ectoplacental Cone migratory
Synciotrophoblast I precursor
Sinusoidal–TGC precursor 1
Sca1+ Trophoblast
Spongiotrophoblast 1
Sinusoidal–TGC precursor 2
Spongiotrophoblast 2
Spiral Artery–TGC
Glycogen Trophoblast
Sinusoidal–TGC
All trophoblasts

**Supplemental Figure 1: Temporal dynamics of *Phlda2* expression across trophoblast populations during placental development**

(a) UMAP projection of *Phlda2* expression single cell RNA sequencing data of mouse placenta. Trophoblast populations are highlighted.
(b) Violin plot of *Phlda2* expression across placenta cell types.
(c) Line graph plotting *Phlda2* expression (y-axis) across developmental timepoints (x-axis) in trophoblast populations, illustrating the temporal regulation and dynamic expression patterns during placental differentiation.

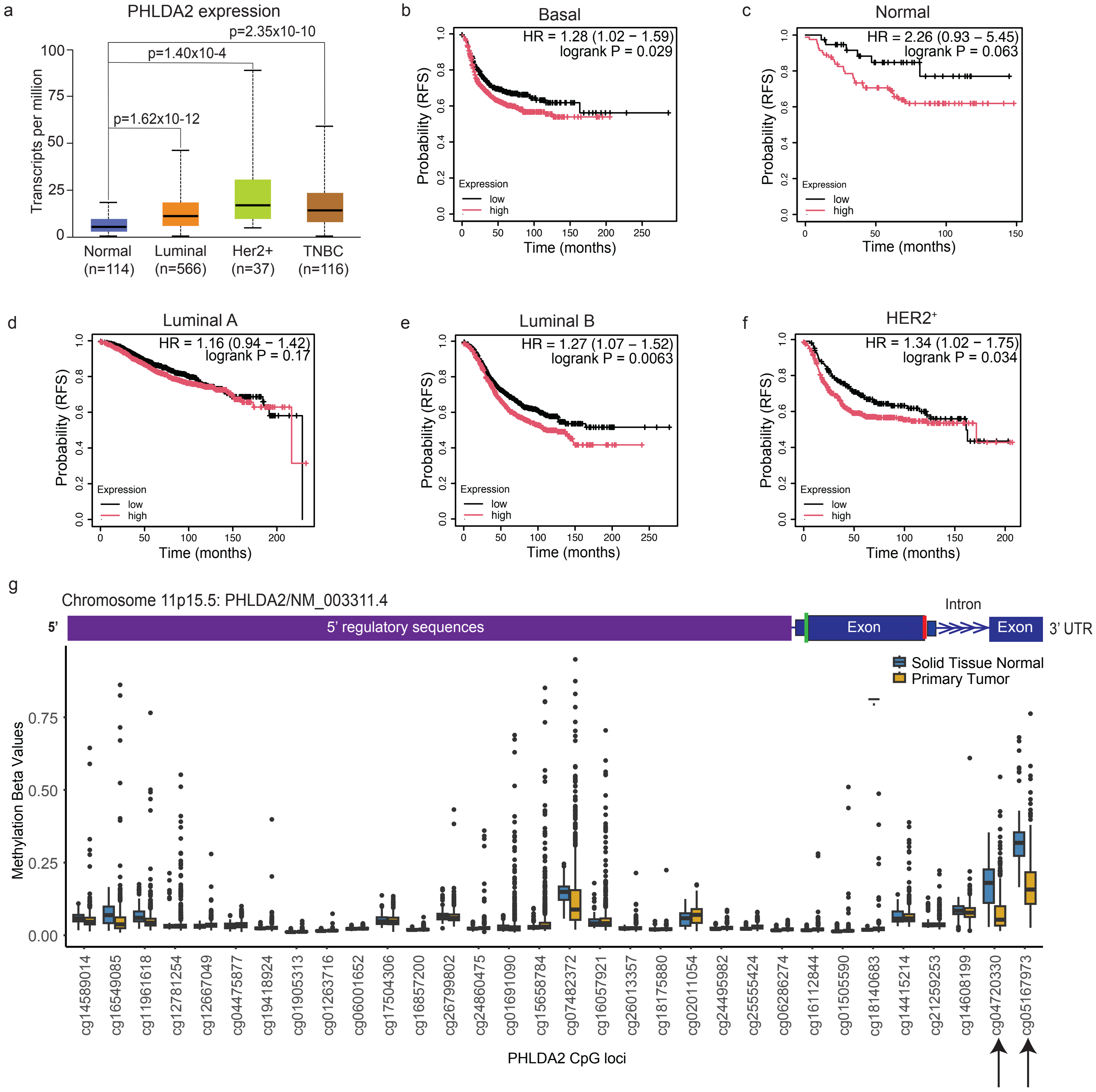
a
PHLDA2 expression
p=2.35x10-10
p=1.40x10-4
p=1.62x10-12
Transcripts per million
Normal (n=114)
Luminal (n=566)
Her2+ (n=37)
TNBC (n=116)
b
Basal
HR = 1.28 (1.02 − 1.59)
logrank P = 0.029
c
Normal
HR = 2.26 (0.93 − 5.45)
logrank P = 0.063
d
Luminal A
HR = 1.16 (0.94 − 1.42)
logrank P = 0.17
e
Luminal B
HR = 1.27 (1.07 − 1.52)
logrank P = 0.0063
f
HER2+
HR = 1.34 (1.02 − 1.75)
logrank P = 0.034
Probability (RFS)
Time (months)
Expression
low
high
g
Chromosome 11p15.5: PHLDA2/NM_003311.4
5'
5' regulatory sequences
Exon
Intron
Exon
3' UTR
Solid Tissue Normal
Primary Tumor
Methylation Beta Values
cg14589014
cg16549085
cg11961618
cg12781254
cg12667049
cg04475877
cg19418924
cg01905313
cg01263716
cg06001652
cg17504306
cg16857200
cg26799802
cg24860475
cg01691090
cg15658784
cg07482372
cg16057921
cg26013357
cg18175880
cg02011054
cg24495982
cg25555424
cg06286274
cg16112844
cg01505590
cg18140683
cg14415214
cg21259253
cg14608199
cg04720330
cg05167973
PHLDA2 CpG loci

**Supplemental Figure 2: *PHLDA2* expression, methylation status, and association with survival and metastasis in breast cancer patients.**

(a) *PHLDA2* expression in healthy breast (n=114) and primary breast tumors (n=1097) representing different breast cancer subtypes Luminal, HER2 positive and Triple Negative. Bar plot shows RNA expression in tissues from the TCGA dataset plotted using the UALCAN tool (ualcan.path.uab.edu). Data are displayed as mean ± SD. *P* value was calculated using a student's t-test.

(b) Kaplan-Meier plots showing probability of RFS in patients with basal subtype breast cancer with high (n=561) vs. low (n=392) expression of *PHLDA2* in their primary tumor tissue. Plots were generated using the KM plotter dataset of mRNA Breast Cancer gene chip and visualized using the KM plotter tool (KMplot.com).

(c) Kaplan-Meier plots showing probability of RFS in patients with Normal subtype breast cancer with high (n=81) vs. low (n=38) expression of *PHLDA2* in their primary tumor tissue. Plots were generated using the KM plotter dataset of mRNA Breast Cancer gene chip and visualized using the KM plotter tool (KMplot.com).

(d) Kaplan-Meier plots showing probability of RFS in patients with Luminal A subtype breast cancer with high (n=957) vs. low (n=852) expression of *PHLDA2* in their primary tumor tissue. Plots were generated using the KM plotter dataset of mRNA Breast Cancer gene chip and visualized using the KM plotter tool (KMplot.com).

(e) Kaplan-Meier plots showing probability of RFS in patients with Luminal B subtype breast cancer with high (n=643) vs. low (n=710) expression of *PHLDA2* in their primary tumor tissue. Plots were generated using the KM plotter dataset of mRNA Breast Cancer gene chip and visualized using the KM plotter tool (KMplot.com).

(f) Kaplan-Meier plots showing probability of RFS in patients with Her2+ subtype breast cancer with high (n=489) vs. low (n=206) expression of *PHLDA2* in their primary tumor tissue. Plots were generated using the KM plotter dataset of mRNA Breast Cancer gene chip and visualized using the KM plotter tool (KMplot.com).

(g) Methylation values for all *PHLDA2* loci in solid normal tissue (n = 139) and tumor tissue (n = 1101). Significantly methylated loci between primary tumor and normal tissue are marked with arrows. Data are displayed as mean ± SD. Diagram of *PHLDA2* gene was generated with UCSC Genome browser and denotes location of loci in the chromosome using the hg37 reference genome.

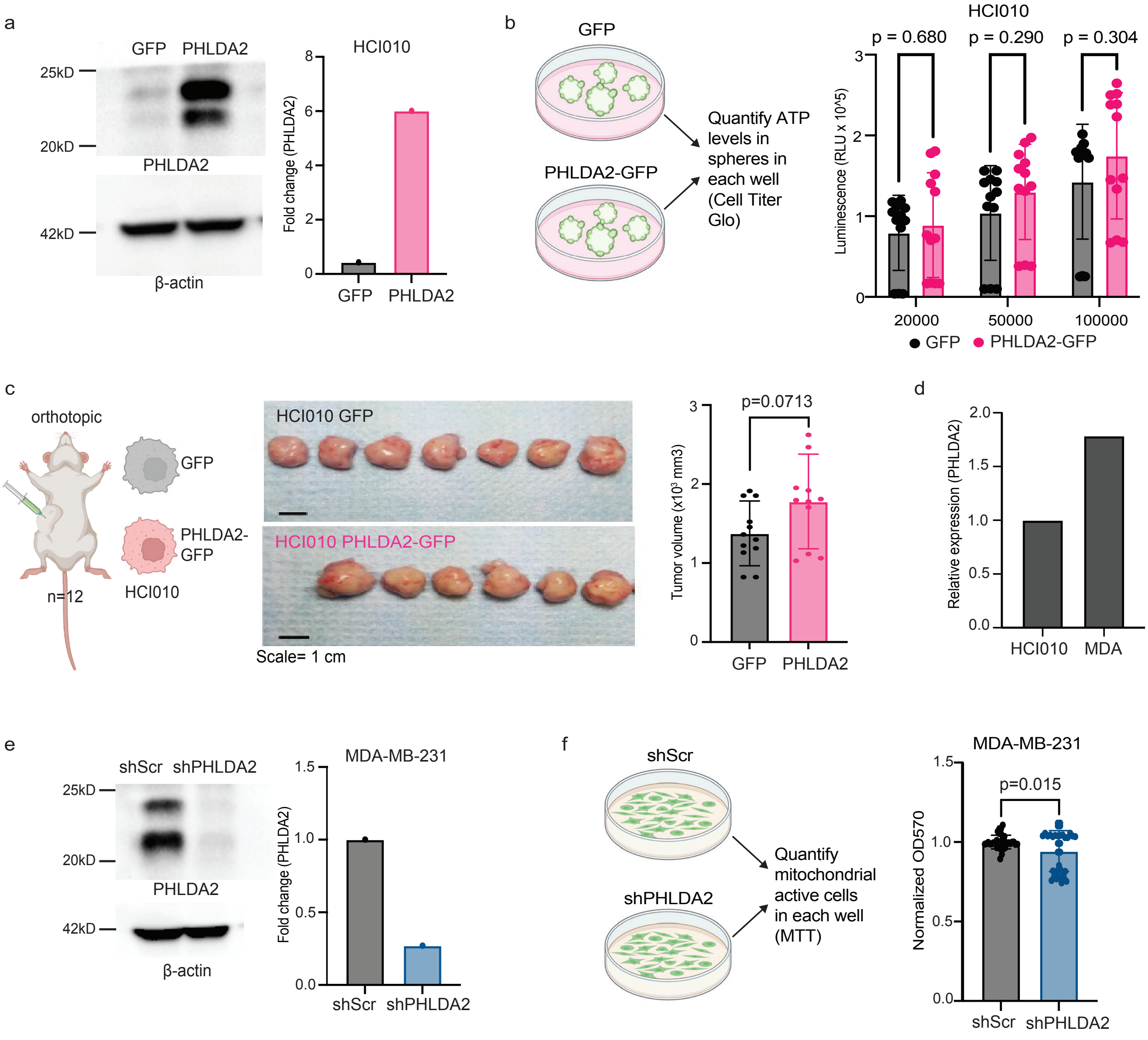

a
GFP PHLDA2
25kD
20kD
PHLDA2
42kD
β-actin
HCI010
Fold change (PHLDA2)
GFP PHLDA2
b
GFP
PHLDA2-GFP
Quantify ATP levels in spheres in each well (Cell Titer Glo)
HCI010
p = 0.680 p = 0.290 p = 0.304
Luminescence (RLU x 10^5)
20000 50000 100000
GFP PHLDA2-GFP
c
orthotopic
GFP
PHLDA2-GFP
n=12
HCI010
HCI010 GFP
HCI010 PHLDA2-GFP
Scale= 1 cm
p=0.0713
Tumor volume (x10³ mm3)
GFP PHLDA2
d
Relative expression (PHLDA2)
HCI010 MDA
e
shScr shPHLDA2
25kD
20kD
PHLDA2
42kD
β-actin
MDA-MB-231
Fold change (PHLDA2)
shScr shPHLDA2
f
shScr
shPHLDA2
Quantify mitochondrial active cells in each well (MTT)
MDA-MB-231
p=0.015
Normalized OD570
shScr shPHLDA2

**Supplemental Figure 3: Modulation of *PHLDA2* expression has limited effects on cancer cell proliferation**

(a) Western blot (left) shows PHLDA2 protein expression in HCI010 GFP (control) and *PHLDA2*-GFP cells that were sorted from primary tumors by flow cytometry. Bar plot (right) shows quantification of PHLDA2 expression normalized to β-actin quantified with densitometry using ImageJ.

(b) Quantification of cell viability and proliferation in organoids generated from HCI010 GFP versus *PHLDA2*-GFP cells by Cell Titre Glo assay. Cells were seeded at three densities (20,000, 50,000, 100,000 cells) (n=3). Bar plot shows luminescence quantification of ATP in each well. P values were calculated using multiple student's t-tests and are represented on the graph.

(c) HCI010 cells were injected into the mammary fat pad of mice to form primary tumors. Images show primary tumors from HCI010-GFP (n=12) and HCI010-*PHLDA2* GFP cells (n=12). Scale bar = 1 cm. Bar plot (right) shows tumor volume ($mm^3$). Data are displayed as mean ± SD. P value was calculated using a student's t-test and is represented on the graph.

(d) Bar graph plotting *PHLDA2* mRNA expression measured by quantitative PCR in HCI010 and MDA-MB-231 cells. Gene expression is normalized to housekeeping gene control, GAPDH, and displayed as fold change relative to HCI010 GFP.

(e) Western blot (left) shows PHLDA2 protein expression in MDA-MB-231 Scrambled GFP (control) and sh*PHLDA2* GFP cells grown *in vitro*. Bar plot (right) shows quantification of PHLDA2 expression normalized to β-actin quantified with densitometry using ImageJ.

(f) Quantification of cell viability and proliferation in MDA-MB-231 Scrambled GFP (control) or sh*PHLDA2* GFP cells by MTT assay. Bar plot represents OD600 which is related to cell proliferation. Each point represents a technical replicate. P value was calculated using a student's t-test and is represented on the graph.

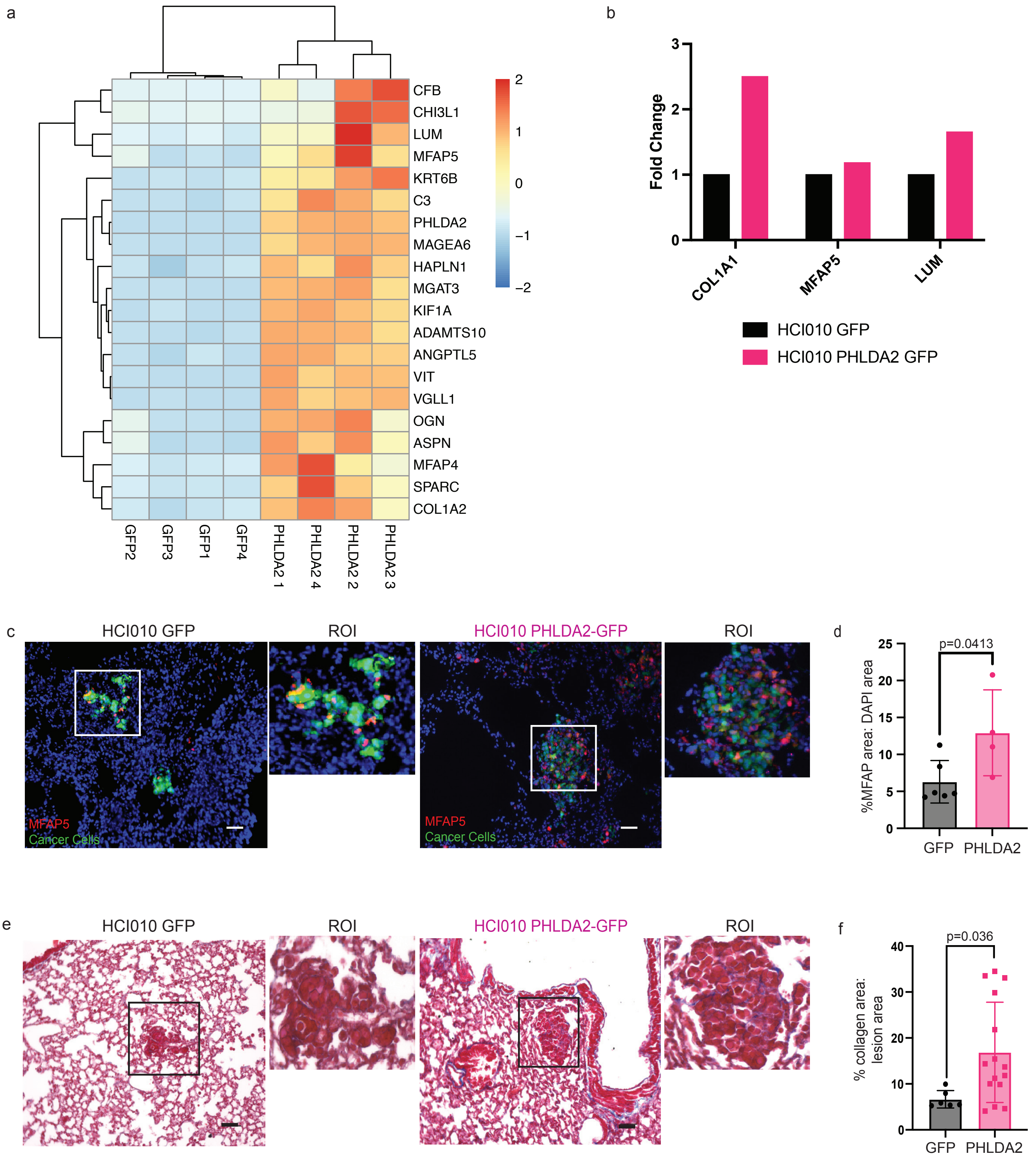

a
CFB
CHI3L1
LUM
MFAP5
KRT6B
C3
PHLDA2
MAGEA6
HAPLN1
MGAT3
KIF1A
ADAMTS10
ANGPTL5
VIT
VGLL1
OGN
ASPN
MFAP4
SPARC
COL1A2
2
1
0
−1
−2
GFP2
GFP3
GFP1
GFP4
PHLDA2 1
PHLDA2 4
PHLDA2 2
PHLDA2 3
b
Fold Change
3
2
1
0
COL1A1
MFAP5
LUM
HCI010 GFP
HCI010 PHLDA2 GFP
c
HCI010 GFP
ROI
HCI010 PHLDA2-GFP
ROI
MFAP5
Cancer Cells
MFAP5
Cancer Cells
d
%MFAP area: DAPI area
25
20
15
10
5
0
p=0.0413
GFP
PHLDA2
e
HCI010 GFP
ROI
HCI010 PHLDA2-GFP
ROI
f
% collagen area: lesion area
40
30
20
10
0
p=0.036
GFP
PHLDA2

**Supplemental Figure 4: *PHLDA2* modulates the ECM in breast cancer lung metastasis**

(a) Heatmap shows top genes 20 genes upregulated in HCI010 *PHLDA2* tumors compared to HCI010 GFP controls.

(b) Bar graph plotting *COL1A1, MFAP,* and *LUM* mRNA expression measured by quantitative PCR in HCI010 *PHLDA2* primary tumors cells (n=1) compared to HCI010 GFP controls (n=1). Gene expression is normalized to housekeeping gene control, GAPDH, and displayed as fold change relative to HCI010 GFP.

(c) Representative images of immunofluorescence staining for MFAP5 (red) with DAPI (blue) and tumor cells (green) in spontaneous lung metastases from mice transplanted with HCI010 GFP and HCI010 *PHLDA2* tumor cells. Scale bar = 50 µm.

(d) Quantification of MFAP5 expression in lung metastasis from (c). Bar graph shows the percent of $MFAP5^+$ signal normalized to $DAPI^+$ area (see Methods). Each point represents one lung (n= 4-6), and represents the average obtained from 5-10 fields (20x). Data are displayed as mean ± SD. P value was calculated using a student's t-test.

(e) Representative brightfield images of collagen deposition (blue) in spontaneous lung metastasis from mice transplanted with HCI010 GFP and HCI010 *PHLDA2* tumor cells. Collagens are visualized stained using Masson's Trichrome staining. Scale bar = 100 µm.

(f) Quantification of collagen density in lung metastasis from (e). Bar graph shows the percent of blue signal area normalized to lesion area (see Methods). Each point represents one lesion (n= 6 HCI010 GFP; n=16 HCI010 PHLDA2). Data are displayed as mean ± SD. *P* value was calculated using a student's t-test.

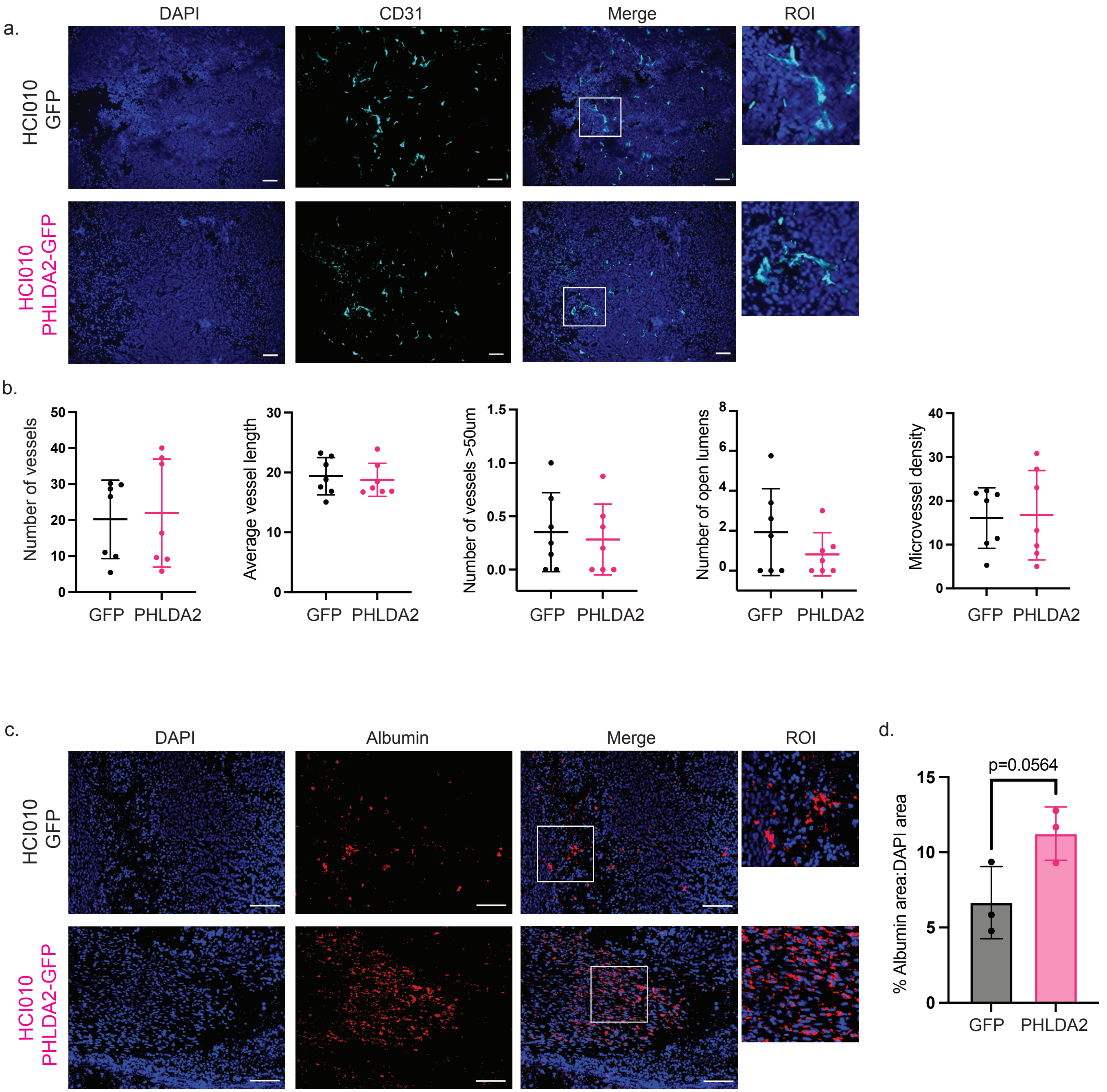

a.
DAPI
CD31
Merge
ROI
HCI010 GFP
HCI010 PHLDA2-GFP
b.
Number of vessels
Average vessel length
Number of vessels >50um
Number of open lumens
Microvessel density
GFP
PHLDA2
c.
DAPI
Albumin
Merge
ROI
HCI010 GFP
HCI010 PHLDA2-GFP
d.
p=0.0564
% Albumin area:DAPI area
GFP
PHLDA2

**Supplemental Figure 5: Increased *PHLDA2* expression does not change vessel morphology but increases vessel permeability in primary tumors**

(a) Representative merged images of immunofluorescence staining for DAPI (blue) and CD31 (cyan) in HCI010 GFP and HCI010 *PHLDA2* primary tumors. Scale bar = 50 µm.

(b) Quantification of vessel morphology parameters including number of vessels (n=7/group), average vessel length (n=7/group), number of open lumens (n=7/group), number of elongated vessels greater than 50µm in length (n=7/group), and microvessel density (n=7/group). See methods for detailed description of quantification. Each point represents one lung tissue value obtained by the average of 5-10 20x objective microscopic fields. Graph is displayed as mean ± SD. *P* values were calculated using student's t-tests, no differences were observed.

(c) Representative merged images of immunofluorescence staining for DAPI (blue) and albumin (red) in HCI010 GFP and HCI010 *PHLDA2* primary tumors. Scale bar = 100 µm.

(d) Quantification of the percent of albumin$^{+}$ area normalized to DAPI$^{+}$ area in HCI010 GFP and HCI010 *PHLDA2* primary tumors (n=3/group). Each point represents one lung tissue value obtained by the average of 5-10 20x objective microscopic fields. Graph is displayed as mean ± SD. *P* values were calculated using student's t-tests.

a

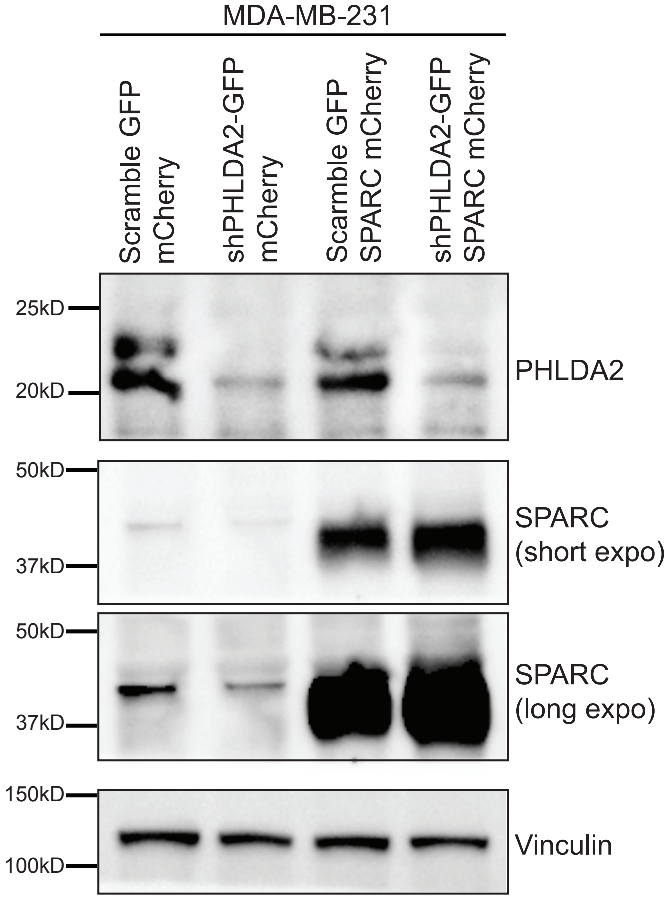
MDA-MB-231
Scramble GFP mCherry
shPHLDA2-GFP mCherry
Scramble GFP SPARC mCherry
shPHLDA2-GFP SPARC mCherry
25kD
20kD
PHLDA2
50kD
37kD
SPARC (short expo)
50kD
37kD
SPARC (long expo)
150kD
100kD
Vinculin

**Supplemental Figure 6: Validation of PHLDA2 knockdown and SPARC overexpression in MDA-MB-231 cells by Western blot**

(a) Western blot shows PHLDA2 and SPARC protein expression in MDA-MB-231 Scramble GFP (control), sh*PHLDA2* GFP, Scramble GFP *SPARC*, and sh*PHLDA2* GFP *SPARC* cells grown *in vitro*. Vinculin is utilized for a loading control.
